# The p*K_a_* values of buried ionizable groups in proteins can be determined by the thermodynamic stability of the protein

**DOI:** 10.64898/2025.11.30.690581

**Authors:** Miranda N. Hurst, Christos M. Kougentakis, Aaron C. Robinson, Bertrand García-Moreno E.

## Abstract

Ionizable groups in hydrophobic environments in proteins usually titrate with anomalous p*Ka* values. The ionization of these buried residues is often coupled to transitions between conformational states. Here we test the hypothesis that because the thermodynamic stability of a protein (ΔG°H2O) determines the probability of conformational transitions, the apparent p*Ka* values of buried residues can be governed by ΔG°H2O. Variants of staphylococcal nuclease (SNase) with either Lys-66 or Lys-92 buried in its hydrophobic interior were engineered along with surface mutations that alter ΔG°H2O without affecting the electrostatic properties of the internal microenvironments of the buried Lys residues. The measured p*Ka* values of these Lys residues largely correlates with ΔG°H2O. NMR spectroscopy was used to demonstrate that the structural changes of the protein backbone coupled to the ionization of Lys-66 or Lys-92 are comparable regardless of the ΔG°H2O of the protein. NMR spectroscopy confirmed that global unfolding of the Lys-92 variant coincides with the apparent p*Ka* of the Lys side chain. The data presented show that the anomalous p*Ka* values measured for internal residues in proteins do not necessarily report on local dielectric or electrostatic properties of the microenvironments around the ionizable group; rather, they can report on the energetics of pH-driven conformational transitions. These data suggest that accurate structure-based calculation of p*Ka* values will require de novo prediction of partially unfolded conformations, and accurate calculation of free energy differences between conformational states, both of which remain formidable challenges.

## Introduction

The hydration of charged groups is one of the strongest forces in biology. For this reason, the charged moieties of ionizable residues in proteins and other biological macromolecules are usually found in water. Ionizable groups that are sequestered from bulk water, or buried in hydrophobic environments, usually signal a specific functional adaptation.^1^ Buried ionizable residues are known to play essential roles in energy transduction processes, including catalysis,^2^ ion^3–5^ and nutrient transport^6,7^, ATP synthesis,^8^ H^+^-coupled electron transfer^9–11^, etc. A detailed molecular and mechanistic understanding of biological energy transduction therefore requires understanding of molecular determinants of the unusual properties of buried ionizable residues.^12^ Ionizable groups in dehydrated, hydrophobic environments in proteins usually titrate with anomalous p*Ka* values that are shifted in the direction that favors the neutral state (i.e., elevated for acidic residues and depressed for basic residues). A systematic study of Lys, Asp, and Glu buried at 25 internal positions in staphylococcal nuclease (SNase) revealed that most of their p*Ka* values are shifted relative to the model p*Ka* values in water, always in the direction that promotes the neutral state, in some cases by as many as 5 pH units. Most Lys, Glu, and Asp residues in internal positions in SNase are neutral at physiological pH.^13–16^

The anomalous p*Ka* values of ionizable residues buried in SNase suggest that the energetic penalty to dehydrate a charged group cannot be sufficiently compensated by the interactions between a buried ionizable moiety and the protein matrix.^17–19^ A large number of crystal structures suggest that the energetics from the shift in p*Ka* is not correlated with the polar or polarizable character of the internal microenvironments of the ionizable moieties.^17,20–22^ Instead, these studies suggest that the p*Ka* values of buried ionizable residues in SNase are determined by conformational transitions coupled to the ionization state of the buried moiety.^16,23–27^ In the case of SNase variants with internal Lys residues, a large number of crystal structures demonstrate that at pH values above their p*Ka*, where the internal Lys residues are neutral, the residues are fully buried in the interior of the protein.^20,27–30^ Characterization by NMR spectroscopy revealed that for 23 of the 25 variants, the ionization of the Lys is coupled to conformational reorganization. The structural reorganization can involve global, local, or partial unfolding, dependent on the position where the Lys is buried.^27,31^ In the reorganized conformations the Lys residue has contact with bulk water. For two variants, V23E and V66K, the locally or partially unfolded conformations have even been observed crystallographically.^32,33^ The evidence that the ionization of buried groups is coupled to structural reorganization suggests an important role for global thermodynamic stability as a determinant of the p*Ka* values of buried ionizable residues in proteins.

The distribution of conformations populated by proteins in solution is governed by their global thermodynamic stability. For proteins with buried ionizable residues, it has been shown that thermodynamic stability decreases at solution pH values above or below the normal p*Ka* value in water for acidic and basic residues, respectively.^19,34,35^ This pH-dependent decrease in stability reflects the anomalous p*Ka* value of these internal group resulting in a plateau in stability once the buried group becomes ionized.^14,36^ Here we examine the hypothesis that, because the ionization of a buried residue is coupled to conformational transitions towards reorganized states where the charge can be hydrated, the p*Ka* values of buried ionizable residues can be influenced by mutations that affect the thermodynamic stability of the protein, which governs the probability of transitions between conformational states.

To test this hypothesis, a family of SNase variants were engineered with surface mutations that increase or decrease thermodynamic stability distal to positions 66 or 92 where the internal Lys were introduced. These two internal positions were chosen for this study because crystal structures show that the ionizable moieties are located in the same general region of the hydrophobic interior^20^, they have similarly depressed p*Ka* values,^14,19^ and their ionization is coupled to conformational reorganization events of different magnitudes.^22,27,31^ Lys-92 titrates with a p*Ka* value of 5.2 and its ionization is coupled to global unfolding,^14,27,28,31^ whereas the ionization of Lys-66 occurs at a p*Ka* value of 5.7 that is coupled to the local unfolding of the C-terminus of helix-1.^19,22,33,36^

Thermodynamic stability was measured with chemical denaturation experiments monitored by intrinsic Trp fluorescence. NMR spectroscopy was used to detect the structural changes of the protein backbone coupled to the ionization of the internal Lys residues. The data showed that the p*Ka* values of buried Lys residues can be shifted towards or away from the normal p*Ka* of Lys in water simply by increasing or decreasing the global thermodynamic stability of the protein through surface mutations that leave the internal microenvironments of the ionizable residues unaltered. Besides contributing mechanistic insight into the molecular determinants of the anomalous p*Ka* values of internal ionizable residues, which has substantial implications for biological energy transduction, these studies present a challenge to structure-based calculations of p*Ka* values. Accurate calculation of p*Ka* values with structure-based methods will require the ability to predict alternative conformational states of proteins, and the ability to calculate with high accuracy the Gibbs free energy of the different conformations. Both remain challenging problems for even the most advanced algorithms developed for conformational sampling and energy calculations.

## MATERIALS & METHODS

### Protein preparation

SNase variants were grown in E. coli BL21/DE3 cells (Invitrogen) that were transformed with a pET24a+ plasmid. Single colonies were selected from a LB agar plate grown overnight at 37°C in the presence of kanamycin. Cells were grown at 37°C in a shaking incubator prior to induction of protein expression and isolation of the proteins from the cells. Proteins were expressed, isolated, and purified as first described by Shortle,^37^ adapted by Byrne et al.,^38^ and detailed by Bolen.^39^ The purified proteins were extensively dialyzed in pure water and flash frozen in liquid nitrogen for storage at -80°C.

### Chemical denaturation experiments

#### Automated titrations

Chemical denaturation experiments were performed for a series of pH values with an ATF-107 automated fluorometer (Aviv Inc.) for both stabilized and destabilized variants, with and without Lys-66 or Lys-92. Buffer and titrant solutions contained 100 mM KCl and 25 mM buffer. The titrant solution additionally contained 6 M guanidinium hydrochloride (GdmCl). The solution pH was adjusted prior to bringing the buffer or titrant to the final volume in a volumetric flask. The buffers used for each range of pH values were: GABA (pH 4-5), potassium acetate (pH 5-6), MES (pH 6-7), HEPES (pH 7-8), TAPS (pH 8-9), CHES (pH 9-10), and CAPS (pH 10-10.5).

Samples were prepared with protein at a concentration of 0.05 mg/mL in the buffer solutions, and a volume of 2 mL was transferred to quartz cuvettes with Hamilton syringes. The intrinsic fluorescence of Trp-140 in SNase variants was monitored at 25 °C, with stirring during measurements. The excitation wavelength was 296 nm, and the emission wavelength was 326 nm. The bandwidth was set to 6.0 nm for both excitation and emission. The ΔG°H2O at each pH value was determined by fitting a two-state unfolding model to the normalized fluorescence signal as described in by Pace.^40^

For the stabilized protein with Lys-92, the stability that was measured at pH 4, below the p*Ka* value, was collected as described above with the following adjustment. Two samples of the protein in buffer were prepared that differ only by the buffer used and the pH of the solution (i.e., at pH 9 and pH 4). The automated titration was collected for the sample at pH 9, and the stability was determined. Without changing the voltage of the PMT, which establishes the intensity of the fluorescent signal for the fully folded protein ensemble in the absence of denaturant at the start of the experiment, the sample at pH 4 was then collected.

### NMR spectroscopy

#### Instrumentation

All data were collected on a Bruker Avance-II 600 MHz instrument equipped with a cryoprobe. Data were processed in NMRpipe^41^ and were assigned and visualized with CCPNMR^42,43^. This study made use of NMRbox: National Center for Biomolecular NMR Data Processing and Analysis, a Biomedical Technology Research Resource (BTRR), which is supported by NIH grant P41GM111135 (NIGMS).^44^

#### Isotopic labelling

Proteins variants of SNase were grown in E. coli BL21/DE3 cells (Invitrogen) that were transformed with a pET24a+ plasmid. Bacteria were grown in minimal M9 media that was inoculated with 5 mL of a LB starter culture grown from a single colony. Uniform isotope labeling of the protein was achieved with the addition of ^15^NH4Cl (1 g/L) and/or ^13^C6-D-glucose (2 g/L). Either a 100 or 250 mL minimal media starter culture was grown to an OD of 0.6 (i.e., optical density from the absorbance measured at 600 nm). The small minimal media culture was used to inoculate a 500 mL or 1 L minimal media cultures, respectively, that were grown to an OD600nm of 0.8-1.0. At the target OD, protein expression was induced with IPTG. The purification steps followed were the same protocol as for SNase proteins that are not isotopically labelled, as described above.

#### NMR sample preparation

Buffers were prepared at a 2x stock concentration that was adjusted to the target pH prior to being brought to a final volume in a volumetric flask. The final concentrations in the NMR sample from the buffer solution were 100 mM KCl and 25 mM buffer for 3D assignment data collection, or 5 mM buffer for spectra collected for the pH titrations. Proteins were concentrated by centrifugation with a pre-washed Amicon Ultra (10 kDa MWCO) centrifugal filter (Millipore Sigma). The protein solution was buffer-exchanged through 3-4 rounds of centrifugation, which involved the addition of up to 4 mL of the 2x buffer stock solution in between spins at 5K rpm for 15 min and 4°C in an Avanti JE centrifuge (Beckman Coulter). The pH of the protein solution was measured after centrifugation to validate buffer exchange was complete. The final sample was made to a total volume of 600 μL, that contained 300 μL of the protein in the 2x buffer solution, 240 or 270 μL Millipore water, and 30 or 60 μL of D2O (Cambridge isotope laboratories, 99.96% deuterated water) for either a 5% or 10% [D2O] sample. The final protein concentrations in each sample was 220-675 μM.

#### NMR pH titrations

For the stabilized and destabilized background proteins, as well as the Lys 66 variants of each, ^15^N-labelled protein samples were made and ^1^H^15^N-HSQC 2D experiments were collected of the amide backbone as a function of pH. The pH titration experiments were performed similarly as previously described by Casteñada et al.^16^ The pH was adjusted with the addition of < 10 μL of 0.1 N HCl or KOH to achieve 0.1-0.2 pH unit steps for the pH titration for the stabilized and destabilized variants with Lys-66. Because the interest was to identify chemical shift perturbations of the protein backbone due to changes in solution pH, the small changes in volume throughout the titration can be ignored. For the stabilized protein with Lys-92 where the nitrogen species is directly detected, the pH was adjusted with the addition of < 1 μL of 1 N HCl or KOH to achieve 0.1-0.2 pH unit steps. The pH of all samples was measured directly in the NMR tube with a 3mm micro-pH electrode (Mettler Toledo).

#### Chemical shift assignments

Doubly-labelled (^15^N/^13^C) protein samples were made to collect 3D assignment experiments for each protein at a pH with optimal ^15^N dispersion where the protein is well folded. Assignments were transferred across spectra collected as a pH titration per variant. For the stabilized background protein, HNCACO, HNCO, CBCA(CO)NH, HNCACB, and HNCA(CO)N 3D experiments^45–48^ were collected for de novo assignment. The (de)stabilized background proteins were compared to spectra previously collected for and assigned for the reference background protein.^16^ Because of the similarity in the chemical shifts between the reference, destabilized, and stabilized background proteins, HNCO and CBCACONH spectra were collected to confirm assignments for the destabilized protein with and without Lys-66 and the stabilized protein with Lys-66.

#### Direct detection of the N^ζ^ species in the stabilized protein with Lys-92

For each pH point, a ^1^H^15^N-HSQC experiment was collected as a 2D spectrum to detect the protein backbone (i.e., ^15^N excitation frequency: 117 ppm, spectral width: 32 ppm). In addition, 1D and 2D ^1^H^15^N-HSQC spectra were collected, separately, that directly observe the amine species of Lys-92 by moving the center of the ^15^N excitation frequency to 26 ppm with a spectral width of 20 ppm. The data was collected as a titration in both directions of pH, starting at low pH and titrating towards the basic range or starting at high pH and titrating towards the acidic range to confirm the pH-induced unfolding was reversible thermodynamically.

## RESULTS

### Stabilized and destabilized background proteins

The crystal structures of SNase variants with V66K or I92K substitutions are shown superimposed in Figure 1 along with the positions that were mutated to increase or decrease thermodynamic stability of the protein. Note that these surface substitutions are spread throughout the protein, far from the ionizable moieties of Lys-66 and Lys-92. Two new background proteins were engineered from the reference protein (Δ+PHS), which has been used previously in studies of p*K*a values of internal Lys in SNase.^14,16,19,36,49^ A stabilized variant of the reference protein was made with three substitutions (T41V/S59A/T82I), and a destabilized variant of the reference protein was made with two substitutions (T33S/A130V). Variants of these background proteins with Lys-66 or Lys-92 were studied as a function of pH as described in the methods.

**Figure 1.**
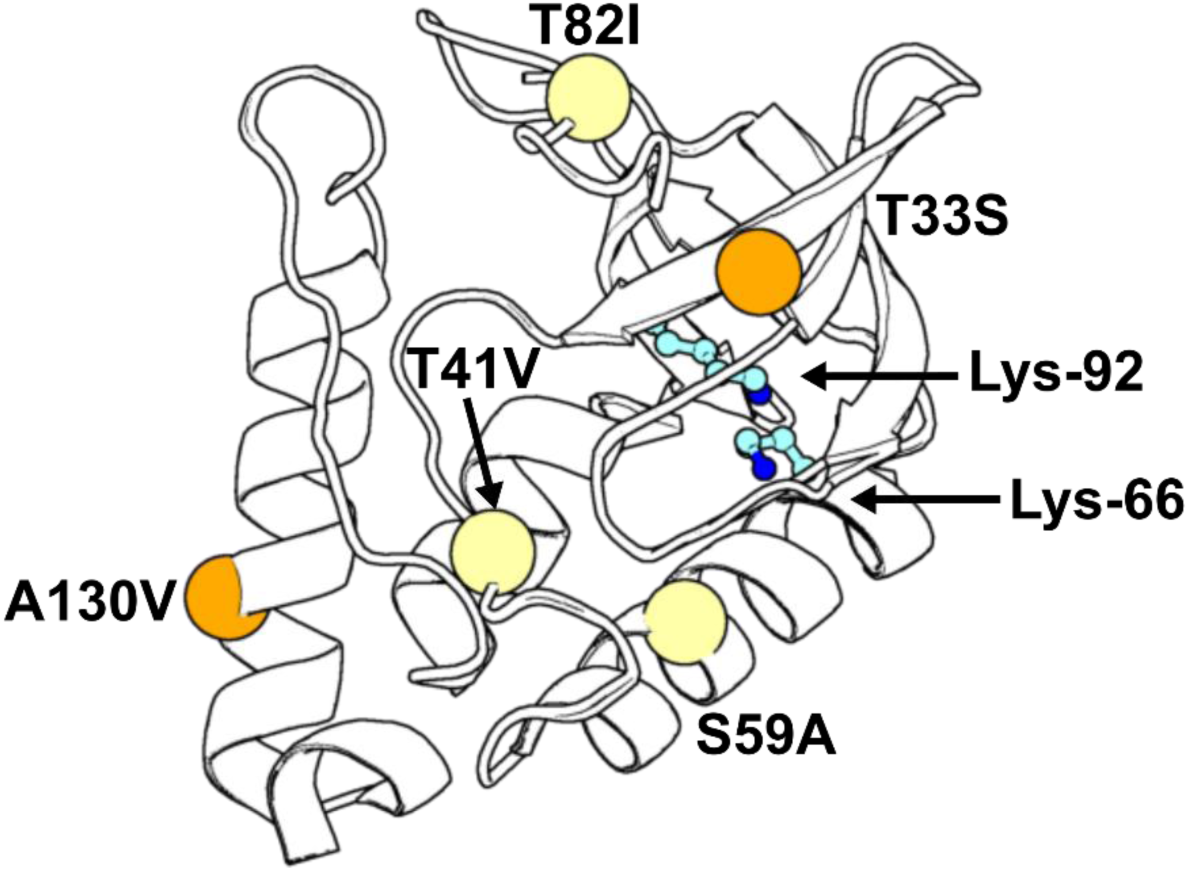
Overlay of the structures woth V66K (PDB: 3HZX) and I92K (PDB: 5E3F) showing the sites of substitutions that increase (yellow) or decrease (orange) the thermodynamic stability of the protein shown. The positions of the side chains of Lys-66 and Lys-92 (cyan, Nζ is blue) are also shown

### Thermodynamic stability of background proteins

The thermodynamic stability of the protein in water (ΔG°H2O) was measured by linear extrapolation of chemical denaturation experiments with guanidinium hydrochloride (GdmCl) (Fig. 2). The unfolding free energy was measured as a function of pH for each protein (Figs. 3 and 4). In the absence of Lys-66 or Lys-92 the thermodynamic stability of the background proteins was pH-independent from pH 5.0 (average) to 9.0. The stabilized background protein has a ΔG°H2O of 13.0 ± 0.1 kcal/mol in the range of pH 4.8-9.0, approximately 1.2 kcal/mol more stable than the Δ+PHS reference protein over the same pH range (Figures 2 and 3). The destabilized background protein has a ΔG°H2O of 9.7 ± 0.1 kcal/mol in the pH range of 5.4-9.0, which is 2.2 kcal/mol less stable than the Δ+PHS reference (Figures 2-3). Fit parameters for all variants are reported in Table 1.

**Figure 2.**
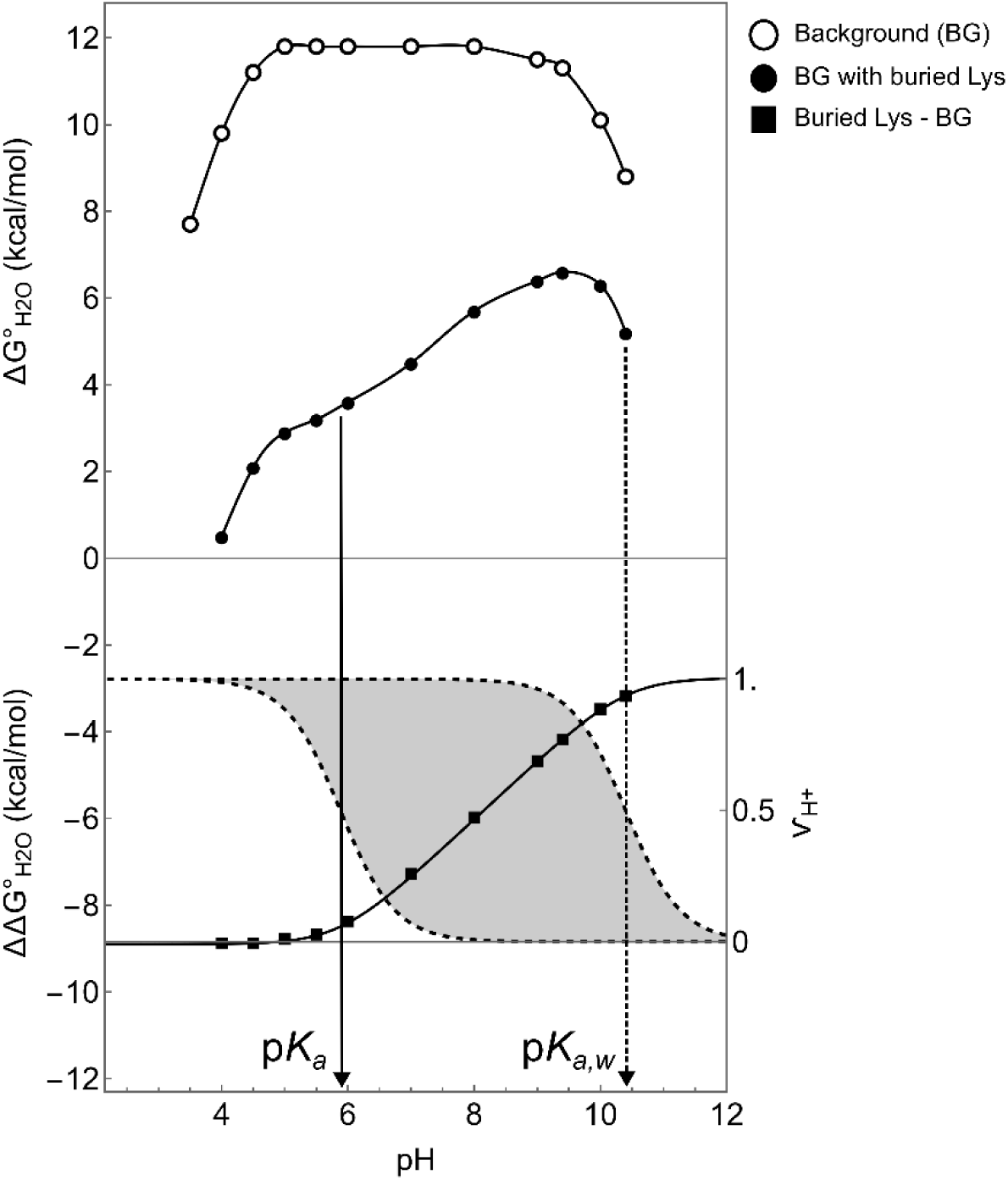
Effect of a buried Lys with a p*Ka* = 6 on the thermodynamic stability (upper panel) and the difference in thermodynamic stability (ΔΔG°H2O) for proteins with and without the buried Lys (lower panel), measured as a function of pH^34^. (**Top panel**) points are the unfolding free energies in water measured at each pH for the background protein (open circles) and of the background protein with internal Lys (solid circles). The lines are to guide the eye. The ΔG°H2O of both proteins decreases at the extremes of pH in the approach to acid and base unfolding. In the physiological range of pH (pH 5-9), the ΔG°H2O is pH-independent for the background protein and highly pH-dependent below the p*Ka* value in water of 10.4 for the variant with internal Lys. (**Bottom panel**) solid squares describe the difference in stability between the two proteins at each pH(variant with Lys minus the background protein). The solid line is a fit to equation 1 to obtain the p*Ka* (solid arrow) with respect to the p*Ka* in water, p*Ka,w* (dashed arrow at pH 10.4). Two H^+^ binding curves (dashed lines, right y-axis) are overlayed in the bottom panel to show that the area between them (shaded region) is equivalent to ΔΔG°H2O(pH).

**Figure 3.**
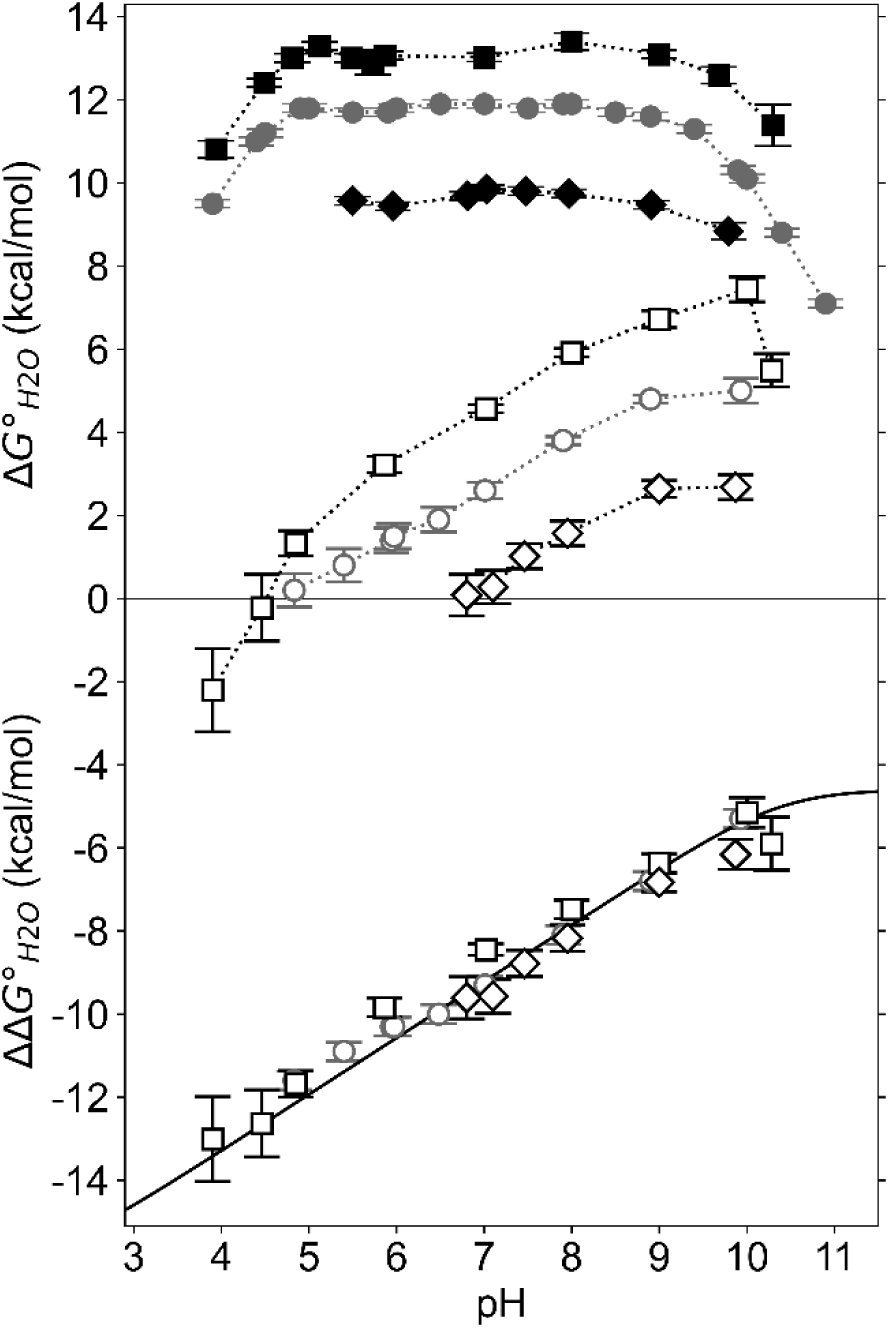
(Top panel) ΔG°H2O as a function of pH for three background proteins with (open symbols) and without Lys-92 (solid symbols). The data for the reference background protein (solid circles) and reference protein with Lys-92 (open circles) were published previously.^14^ The lines are to guide the eye. Data for the stabilized proteins are shown as squares and for destabilized proteins are shown as diamonds. **(Bottom panel)** The data reflect the difference in curves in the top panel. The solid line is a simulation of equation 1, where ΔΔGmut was set to -4.6 kcal/mol (the value in the reference I92K protein), p*Ka,w* was set to the normal p*Ka* value in water for Lys of 10.4, and p*Ka* was artificially set to 0.

**Figure 4.**
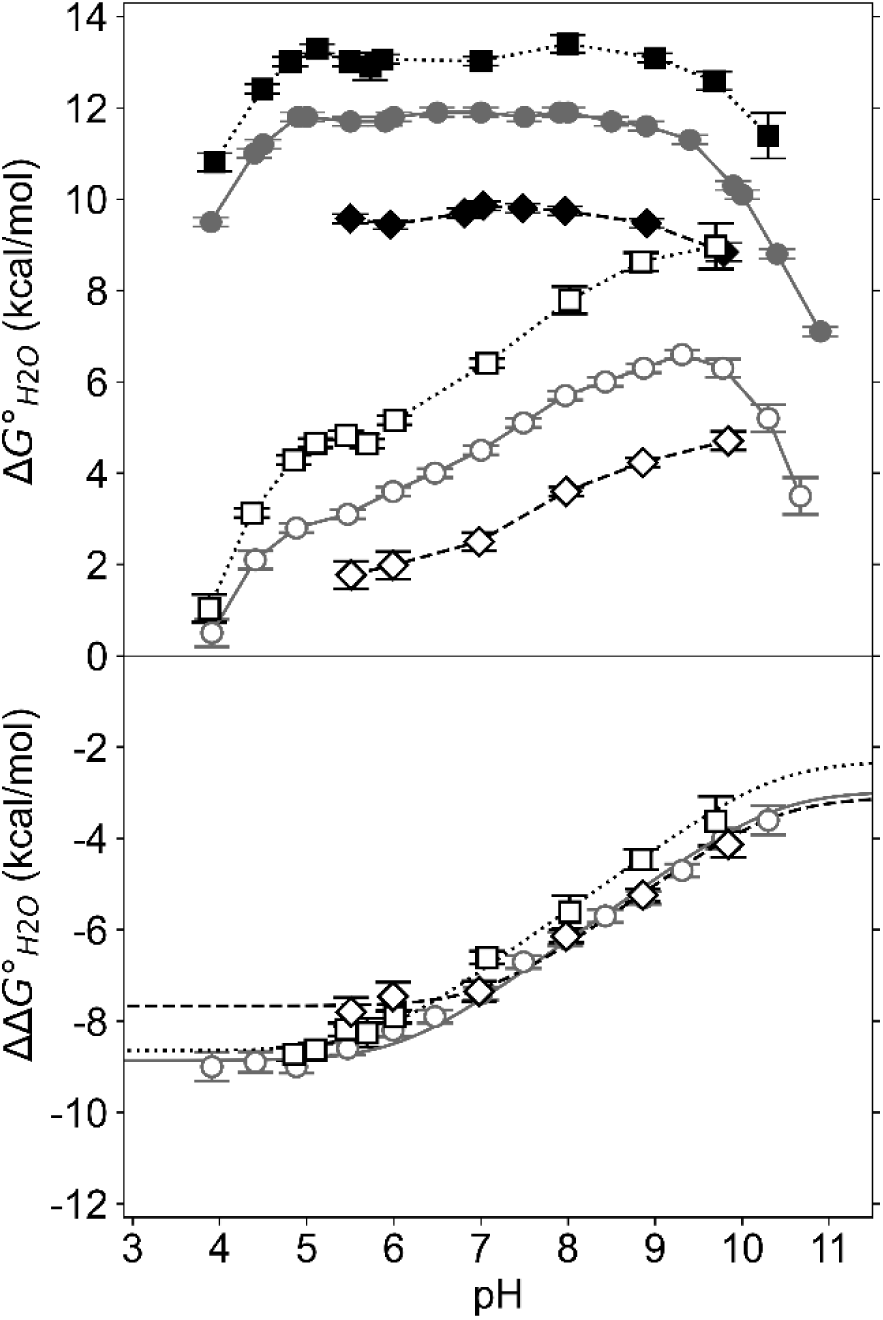
(Top panel) ΔG°H2O as a function of pH for three background proteins with (open symbols) and without Lys-66 (solid symbols). The data for the reference background protein (solid circles) and reference protein with Lys-66 (open circles) were published previously.^14^ The lines are to guide the eye. Data for the stabilized proteins are shown as squares and for destabilized proteins are shown as diamonds. **(Bottom panel)** The data reflect the difference in curves in the top panel. The solid line is a simulation of equation 1, where ΔΔGmut was set to -4.6 kcal/mol (the value in the reference V66K protein), p*Ka,w* was set to the normal p*Ka* value in water for Lys of 10.4, and p*Ka* was set to the fit value in Table 1.

**Table 1.**
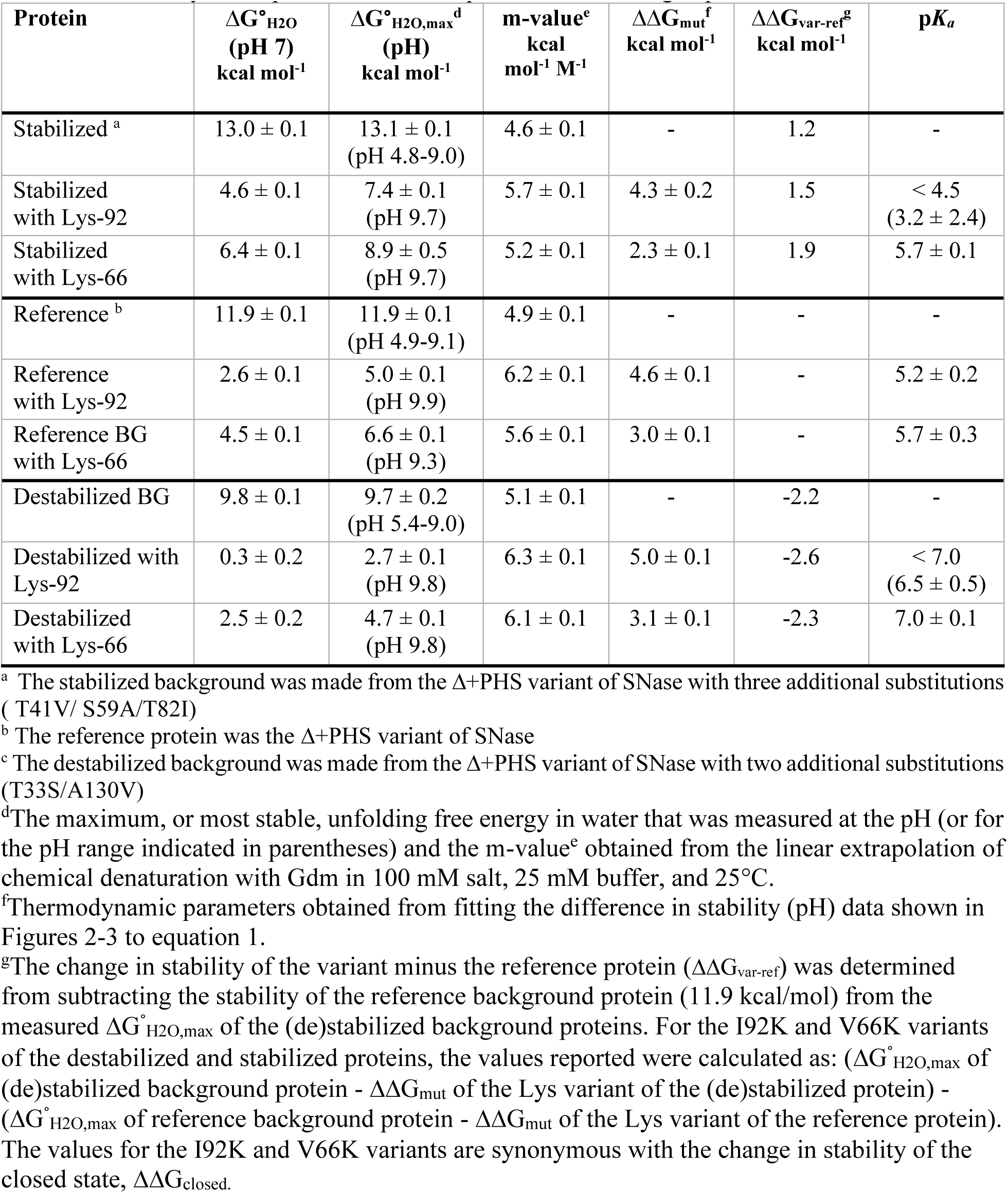
Thermodynamic parameters from equilibrium unfolding experiments

### Thermodynamic stability of background proteins with internal Lys residues

The apparent p*Ka* of the buried Lys residues were determined by fitting Eq. 1 to the pH-dependent difference in stability of proteins with and without the internal Lys:

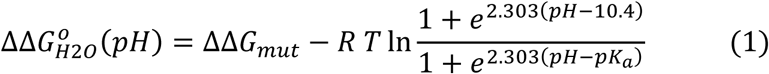

The p*Ka* of Lys in water is fixed as 10.4. The non-electrostatic component of the consequences of the mutation on stability (ΔΔGmut) can be estimated at basic pH values at or above the normal p*K*a of Lys in water. Below the normal p*Ka* of Lys in water, the ΔG°H2O decreases by 1.36 kcal/mol per pH unit as a result of the depressed p*K*a value of the internal Lys. All variants with internal Lys residues studied here exhibited this characteristic pH dependence observed previously in variants of SNase with internal Lys with depressed p*K*a values.^14,19,36^ This is illustrated in the simulation shown in Fig. 2 that shows the relationship between the shape of the stability curves measured as a function of pH (upper panel), the shape of the difference in stability measured in proteins with or without the internal Lys residues (lower panel), and the shift between the p*Ka* value for Lys in water (p*Ka,w*) and the buried state (p*Ka*). The pH sensitivity levels off at pH values below the p*K*a value where the Lys is fully ionized as the result of the protein either globally unfolding or the protein adopting an alternative conformational state in which the Lys is charged and exposed to bulk water. At pH values below the p*Ka*, the Lys residue is fully charged across all proteins in solution.

### p*Ka* of Lys-92 from equilibrium thermodynamic data

The thermodynamic stability of the stabilized and destabilized proteins, with and without Lys-92, are shown as a function of pH in Figure 3. The effect of the stabilizing and destabilizing mutations can be observed on the variants without Lys-92, in the displacements along the y-axis (Fig. 3 upper panel). The slope of ΔG°H2O vs pH is similar in the three variants with Lys-92, and consistent with what is expected from a Lys with a depressed p*Ka*. Note that the pH-dependent stability for variants with Lys-92 continued to decrease until the protein underwent acid-induced unfolding (Fig. 3 upper panel). For the destabilized variant with Lys-92, the midpoint of acid unfolding occurs at pH 7, approximately 2 pH units higher than the midpoint of the reference protein (pHmid = 5.2). In contrast, the stabilized variant with Lys-92 has an acid unfolding midpoint that has shifted to pH 4.5, below that of the reference protein. The p*Ka* values for Lys-92 obtained from the fit of equation 1 to the difference between the stability curves (Fig. 3 lower panel) are 3.2 ± 2.4 in the stabilized protein and 6.5 ± 0.5 in the destabilized protein (Table 1). The large error in the p*Ka* for Lys-92 in these proteins is due to an insufficient number of points in the baseline of the ΔΔG°H2O (pH) curve below the p*Ka* for fitting. For the stabilized protein with Lys-92, ΔG°H2O was measured at pH 4, below the acid unfolding midpoint of 4.5, in an attempt to increase the number of pH points in the unfolded state to improve the error in the fit for p*Ka*. As described in the Methods, at pH 4 the destabilized variant with Lys-92 exhibited a large initial decrease in fluorescent signal compared to samples at higher pH suggesting that even in the absence of denaturant there is a significant population of unfolded protein. The fluorescent signal in the native state was fixed at 8x the intensity of the denatured baseline consistent with what was observed previously for many variants of SNase.^50^ A similar value was obtained when the native baseline was fixed relative to the sample in the absence of denaturant at pH 9.

### p*Ka* of Lys-66 from equilibrium thermodynamic data

The behavior of the variants with Lys-66 differs from that of the variants with Lys-92 but is consistent with the behavior observed for the majority of other SNase variants with internal Lys studied previously^14,31^ and illustrated in Fig. 2. The pH-dependent ΔG°H2O of the proteins with and without buried Lys-66 are shown in Figure 4. For each protein with Lys-66, the pH-dependence of stability plateaus prior to the onset of acid unfolding (Fig. 4 upper panel), suggesting that Lys-66 becomes fully charged by a different mechanism than observed for Lys-92. The p*Ka* values of Lys-66 are 5.7, 5.7, and 7.0 in the reference, stabilized, and destabilized proteins, respectively (Table 1). In all three variants, the combined energetic cost to introduce Lys-66 in the neutral state and shift the p*K*a is ∼8 kcal/mol relative to the stability of the background proteins. Prior to acid unfolding, ΔG°H2O for the Lys-66 variants range between 2 and 5 kcal/mol, suggesting each variant remains in a predominantly folded state, albeit one where Lys-66 is ionized.

### p*Ka* values from NMR spectroscopy

To further corroborate the apparent p*K*a values measured with equilibrium thermodynamic methods, an attempt was made to measure these values directly by NMR spectroscopy. For the variants containing Lys-92, the side chain amino group was detected in the stabilized protein with a traditional ^1^H^15^N-HSQC experiment by moving the ^15^N carrier frequency to 26 ppm which is rare, and quite surprising.^30,51,52^ Figure 5B shows two peaks for Lys-92 at pH 5.8: a major peak with chemical shifts of 0.46 ppm in ^1^H and 21.2 ppm in ^15^N, and a minor peak chemical shifts of 0.43 ppm in ^1^H and 20.4 ppm in ^15^N. Both the relative intensity and position of the peaks are consistent with the NH2 and NHD species of a neutral Lys given the ratio of 90% H2O: 10% D2O used for the 2D experiments. The NHD species was confirmed by the disappearance of this peak in a coaxial NMR tube, where the NMR sample is not in contact with the D2O needed for locking.^52^ The fact that the NH2 and NHD peaks were detectable by this experiment type indicates unambiguously that the neutral species of Lys-92 undergoes significantly slower dynamics than a typical Lys exposed to bulk solvent. Furthermore, the extreme upfield shift of the H^ζ^ indicates that the local environment is highly shielded, consistent with the moiety being buried inside the protein.^30^ For comparison, a peak consistent with a charged, solvent exposed NH3^+^ species with chemical shifts at 32.6 ppm ^15^N and 7.45 ppm ^1^H,^30,51,52^ is shown in Fig. 5A. Below pH 4.4, where the protein undergoes acid-induced unfolding this is the only peak detectable.

**Figure 5.**
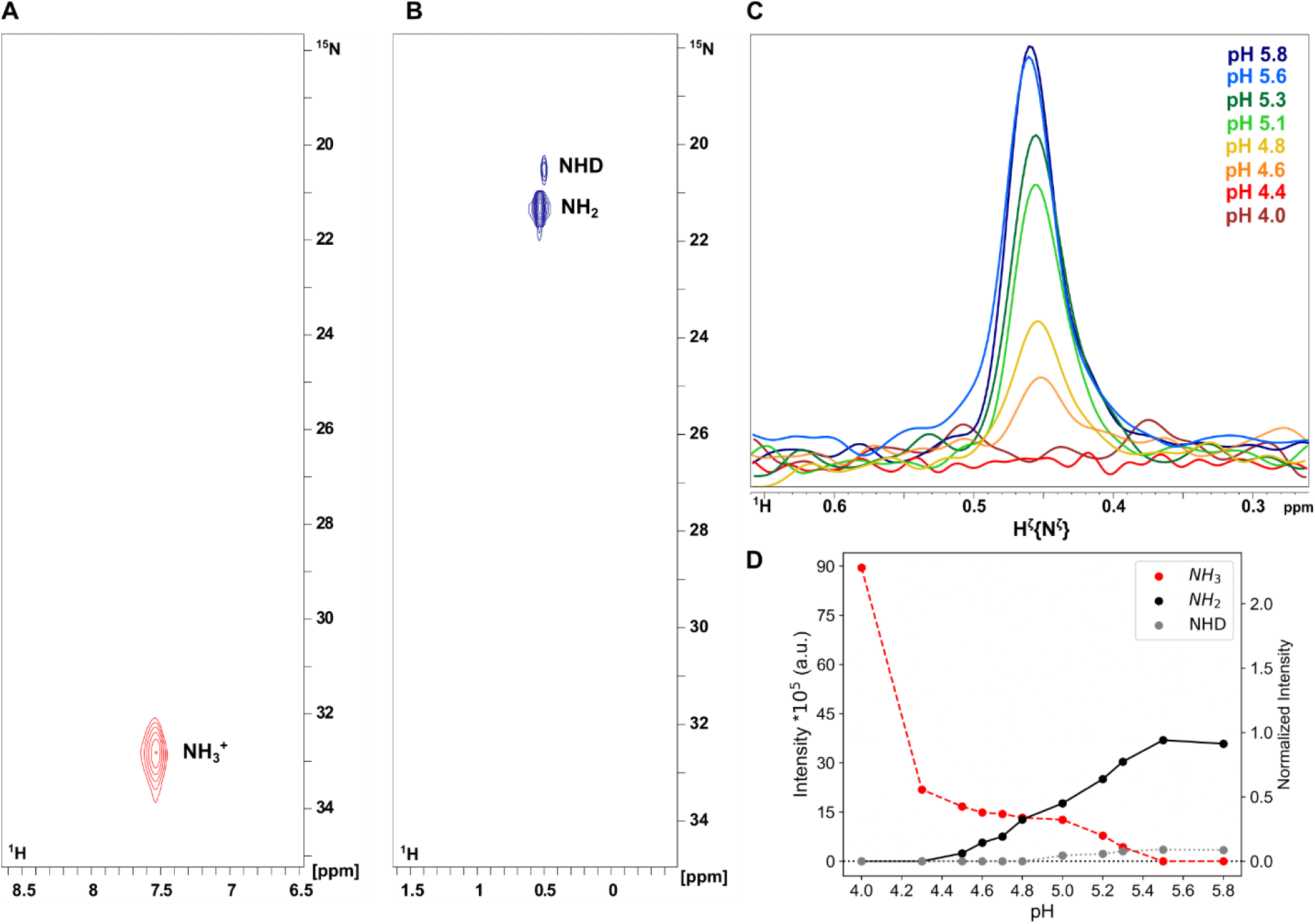
Direct detection of the Hζ{Nζ} of Lys-92 in the stabilized protein observed by an ^1^H^15^N-HSQC with the ^15^N excitation frequency moved to the range of Lys side chains (21-32 ppm) at 25°C. **(A)** A 2D spectrum of a charged NH3^+^ species that is indistinguishable from other solvent exposed, charged Lys in the protein. **(B)** A 2D spectrum of the neutral NH2 and NHD species of Lys-92 in the stabilized variant detected above the p*Ka* value. **(C)** 1D experiments following the Hζ peak intensity of the NH2 species in the presence of 5% D2O as a pH titration. **(D)** The peak intensities from the 2D spectra plotted as a function of pH for the charged species (NH3^+^; red, dashed line), the neutral species (NH2; black, solid line), and the neutral NHD species observed in the presence of 10% D2O (gray, dotted line). The normalized intensity on the right y-axis was calculated relative to the total intensity of the three species at pH 5.8. The lines are to guide the eye.

Figure 5C shows a series of 1D ^1^H^ζ,15^N^ζ^-HSQC experiments following the H^ζ^ peak intensity over the course of a pH titration. The disappearance of the H^ζ^ peak at pH 4.4 is consistent with a p*Ka* of <4.5 for Lys-92 in the stabilized protein determined by analysis of ΔΔG°H2O (pH). However, in these NMR experiments the ratio of charged:neutral Lys species has not been determined explicitly. Rather, these data support that the protein acid unfolds near pH 4.4, rendering the measurement of any intrinsic p*Ka* value for Lys-92 impossible. Figure 5D shows the intensities of the NH3, NH2, and NHD species observed in 2D spectra plotted as a function of pH. The peak pertaining to the charged Lys-92 species is undetectable above pH 5.4, only starting to appear near the pH of acid unfolding of Lys-92. While the peak for the charged species increases in intensity as the peak for the neutral species decreases in intensity, its signal at pH 4 has increased in intensity by 2.5x relative to the signal normalized to all three species at pH 5.8. The peak was not further interpreted because the chemical shifts of the charged species are indistinguishable from the signal of the surface Lys residues natural to the protein.

An attempt was made to detect Lys-66 given that the crystal structure suggests its side amino moiety exists in approximately the same microenvironment and would be expected to have a similar chemical shift. Despite repeating the same procedure over a range of temperature from 5-25°C, neither the charged nor the neutral species was detectable by ^1^H^15^N-HSQC.

### Structural reorganization coupled to the ionization of Lys-92

NMR spectroscopy was also used to examine the coupling between H^+^-binding to the internal Lys-92 and structural reorganization of the protein backbone with the goal of identifying the residues affected by the ionization of the buried Lys to compare in proteins with different thermodynamic stabilities. Fig. 6A-D shows 2D ^1^H^15^N-HSQC experiments of the backbone amides for the stabilized protein with Lys-92 between pH 4.3 and 7.0, both above and below the apparent p*K*a of Lys-92. The spectrum of the amide backbone at pH 7 shows a pattern consistent with the background protein suggesting the native, folded conformation is the dominant species in solution above the p*Ka* of Lys-92 in the stabilized protein. At pH 5.3, near the p*Ka* of Lys-92 in the reference protein, the folded conformation is still largely populated, though there is some evidence of slow exchange with an unfolded state. By pH 4.6, just above the observed p*Ka* of Lys-92, there is a dramatic shift in populations with the globally unfolded state becoming the dominant species as evidenced from the uniform decrease in intensity of peaks pertaining to the folded state. Just below the p*Ka* at pH 4.3, the ensemble has completely transitioned into a globally unfolded state. The uniform loss in intensity of resonances corresponding to the folded state near pH 4.5 without line broadening suggest the protein undergoes a two-state transition between the fully folded and the fully unfolded states without every populating a partially unfolded state in which the ionizable moiety of Lys-92 can be protonated and exposed to bulk water.

**Figure 6.**
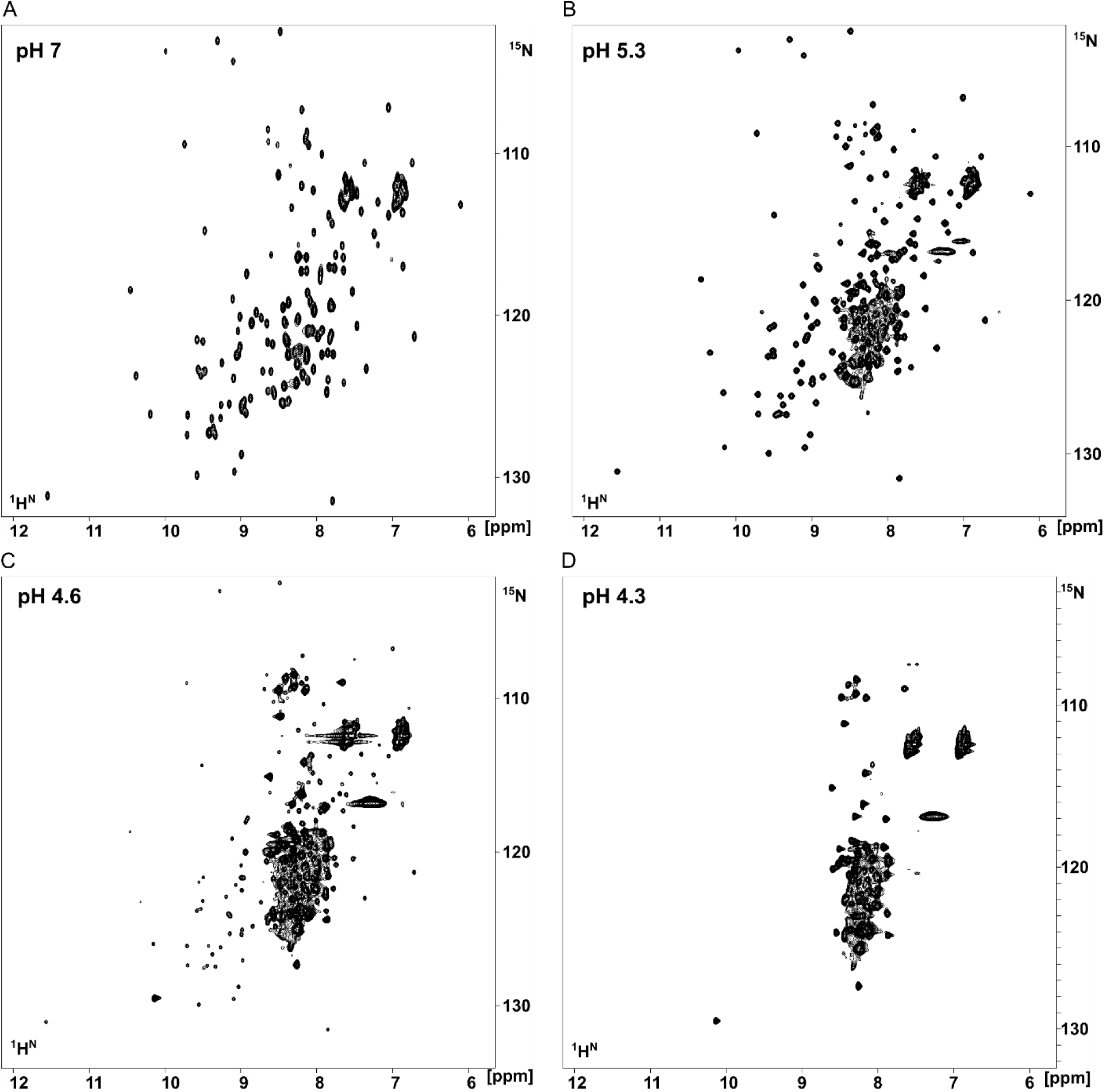
^1^H^15^N-HSQC spectra of the amide backbone of the stabilized protein with Lys-92 at pH 7.0 **(A)**, pH 5.3 **(B)**, pH 4.6 **(C)**, and pH 4.3 **(D)**. The data show that the proteins enter slow exchange between folded and globally unfolded populations around 1 pH unit of the p*Ka* value of < 4.5 for Lys-92. An overlay of the spectra at pH 5.3 and 4.3 is shown in the supporting information.

### Structural reorganization coupled to the ionization of Lys-66

^1^H^15^N-HSQC spectra of the backbone amides were collected as a function of pH for the stabilized and destabilized proteins with and without Lys-66 (see supporting information). The combined HN chemical shift perturbations (CSPs) were calculated at pH values above and below the p*Ka* value of Lys-66 in stabilized and destabilized proteins with and without Lys-66 (supporting information). For the stabilized protein with Lys-66, CSPs were calculated between pH 7.1 and 5.0, 1.4 pH units above and 0.7 pH units below the p*Ka* value of 5.7, respectively. For the destabilized protein with Lys-66, CSPs were calculated between pH 8.0 and 6.3, 1 pH unit above and 0.7 pH units below the p*Ka* value of 7.0. The chemical shifts from spectra collected at pH 8.0 were used instead of pH 8.5 for CSP calculations because the chemical shifts at pH 8.0 and pH 8.5 were similar for the destabilized protein with Lys-66, but for the destabilized background protein, several residues lose resonance at pH 8.5 as the pH approaches the pI of the protein (9.1). For both the stabilized and destabilized proteins with Lys-66, the change in the chemical shift perturbations (ΔCSPs) due to the ionization of Lys-66 was calculated by subtracting the CSPs of the background protein from the CSPs of the Lys-66 variant. The ΔCSPs are shown mapped onto the crystal structure of the reference protein with Lys-66 in Figure 7.

**Figure 7.**
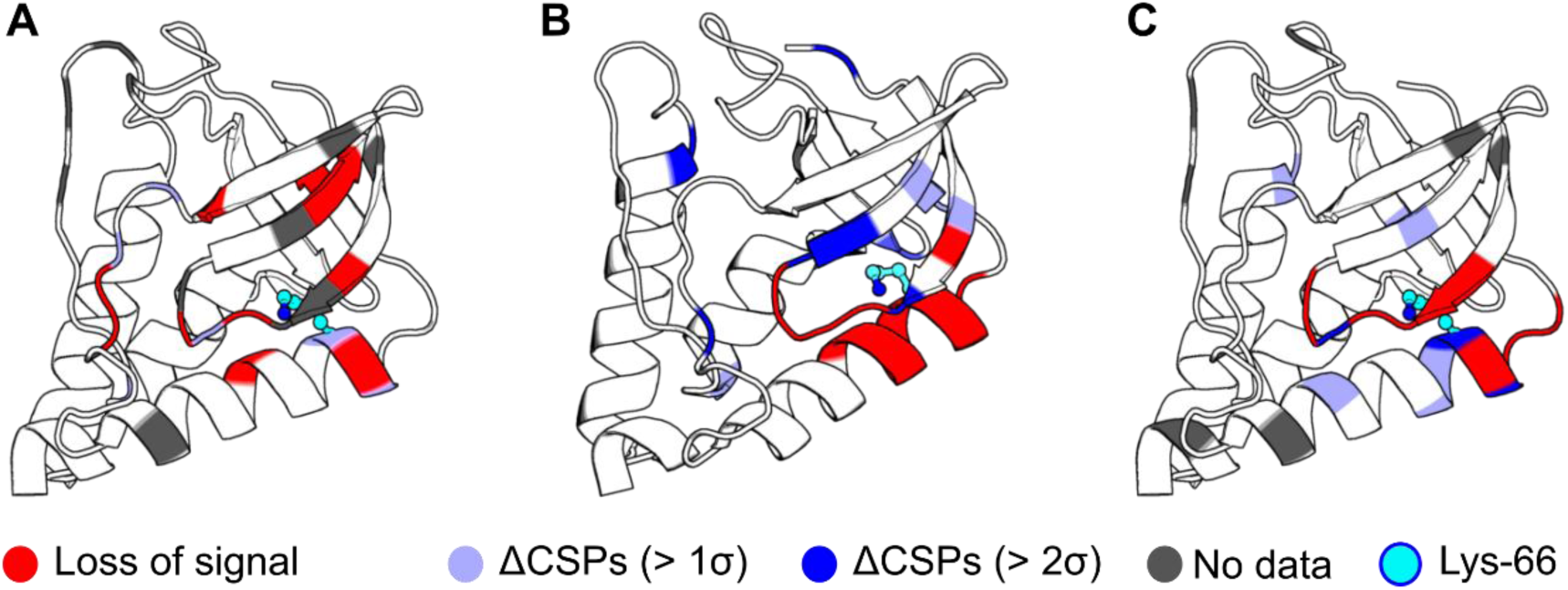
Residues that are affected by the ionization of buried Lys 66 in the **(A)** destabilized, **(B)** reference, and **(C)** stabilized proteins determined by NMR spectroscopy mapped onto the crystal structure of the reference protein with V66K^17,29^. In **(A)** and **(C)**, the combined HN CSPs were calculated from ^1^H^15^N-HSQC spectra collected at pH above and below the p*Ka* value of Lys-66 (see supporting information). In **(B)**, the combined H,N,Cα,Cβ CSPs were calculated from 3D HNCACB experiments collected above and below the p*Ka* value of Lys-66.^31^

Also shown in Figure 7 are the ΔCSPs determined for the reference protein with Lys-66, considering the chemical shifts of the H^N^,N^H^,C^α^,C^β^ ΔCSPs determined from 3D HNCACB experiments collected at pH values above and below the p*Ka* value. The significance of reporting ΔCSPs is to remove from consideration residues that exhibit the same pH-dependence in both the background and Lys-66 proteins or residues whose CSPs are insignificant after consideration of random fluctuations in the background protein. Because the loss of signal in ^1^H^15^N-HSQC spectra can be caused by changes in the ^1^H exchange rate with water, 2D (HA)CACO experiments were collected as a function of pH to directly detect ^13^C for the stabilized protein with and without Lys-66 (supporting information). The combined CaCO CSPs were calculated at the same pH as described for ^1^H^15^N (supporting information). For both datasets, the same region of the protein is affected, as shown by comparing the (Δ)CSPs for HN and CaCO nuclei for the stabilized protein with and without Lys-66 (supporting information). For all of the Lys-66 variants, structural perturbations were localized to the C-terminus of helix-1 and strands β1 and β2, in direct proximity to the side chain of Lys-66. Previous studies of Lys-66 variants show the same local response to ionization of the side chain without major reorganization of the rest of the protein.^22,33^

## DISCUSSION

Transitions between conformational states of a protein are pH-sensitive when ionizable residues have different p*K*a values in the different conformational states.^53^ The energetic contribution from the shift in p*K*a is depicted in Figure 8, which are the same energetics measured in Figure 2. When buried in the hydrophobic interior of the reference SNase protein, Lys-92 and Lys-66 titrate with anomalous p*Ka* values of 5.2 and 5.7, respectively. Relative to the normal p*K*a of 10.4 for Lys in water, these Δp*Ka* represent a ΔG°H2O of at least 6.4 kcal/mol. Not surprisingly, given that this ΔG°H2O is comparable to the global stability of SNase, the ionization of these residues is coupled to conformational transitions. These two cases illustrate why ionizable residues are frequently used as triggers of conformational transitions in proteins: a single Lys buried in a hydrophobic environment of a protein can destabilize the protein by a factor comparable to the average ΔG°H2O of globular proteins, 5-15 kcal/mol.^54,55^ The hydration free energy of a single charged atom can exceed the total stability of many globular proteins. This is the reason proteins evolved such that their ionizable moieties are always exposed to water except in special situations where, for functional purposes, they are sequestered in dehydrated environments. Buried in the neutral state in these environments, these residues become triggers that can drive conformational events in response to changes in pH.

**Figure 8.**
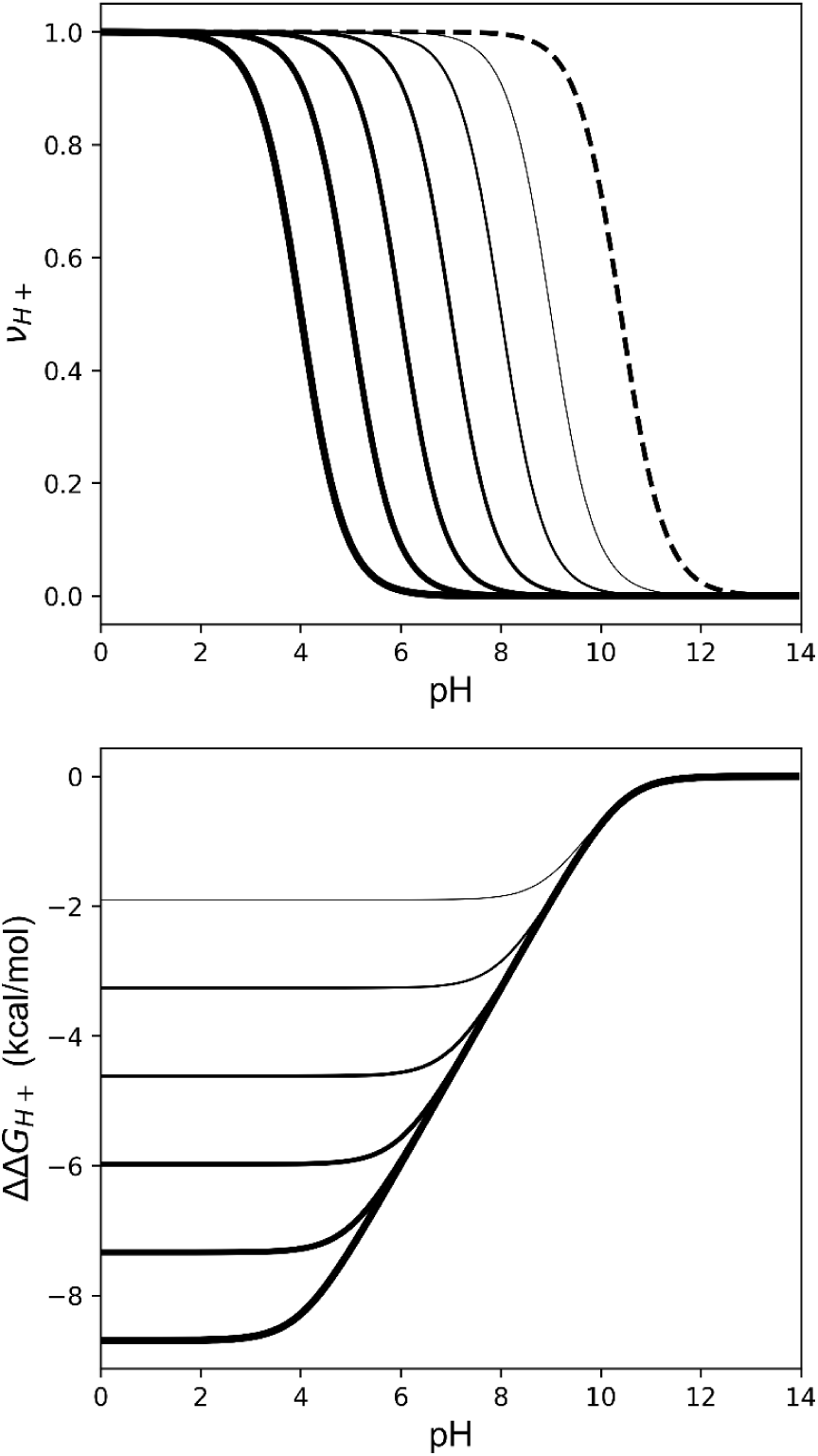
Simulations of the free energy of H^+^ binding due by a single Lys residue with shifted p*Ka* value. **(Top)** Simulated H^+^ titration curves for a Lys with a normal p*Ka* value in water (dashed line, pH 10.4) and with hypothetical p*Ka* values of 9, 8, 7, 6, 5, 4 (solid lines). **(Bottom)** Simulation of the difference in free energy of H^+^ binding due to the shift in the p*Ka* value from water for p*Ka* values 9 to 4 (thinner to thicker solid lines). The free energy was obtained by integrating the area between H^+^ titration curves shown in solid lines and the reference dashed curve for the p*Ka* in water of 10.4 (at 298 K; ΔΔ𝐺_𝐻+_ = −2.303 𝑅 𝑇 ∫(𝜈_𝐻+,1_ − 𝜈_𝐻+,2_) 𝑑𝑝𝐻).

Here we have tested the idea that the p*Ka* values of internal ionizable groups are not determined by the local electrostatic or dielectric properties around the ionizable moiety, but rather, by the global stability of the protein because that determines the propensity of the protein to transition into conformational states where the buried group can be hydrated when charged. This implies that the p*Ka* values of internal ionizable groups can be tuned by changes to the thermodynamic stability of the protein. The fact that the apparent p*Ka* values of Lys-66 and Lys-92 could be altered by secondary mutations far from the local microenvironment of the ionizable moiety that either increased or decreased the global stability of the protein demonstrates that the thermodynamic stability of the protein can be an important determinant of the p*Ka* values of buried groups.

The case of Lys-92 is illustrative. Its apparent p*Ka* values in the stabilized, reference, and destabilized variants are 6.5, 5.2 and < 4.5 (Table 1), consistent with the idea that the apparent p*Ka* value of Lys-92 can be altered simply by changing the global thermodynamic stability of the background protein. The true p*K*a of Lys-92 in the buried state can never be measured, not even in stabilized forms of the protein, because the protein unfolds whenever the buried Lys residue becomes ionized. For these variants with Lys-92, the globally unfolded state appears to be the lowest energy conformation where the amino moiety of Lys-92 can be solvated. The substantial depression in the p*K*a means that a large amount of ΔG°H2O is lost as the pH decreases until the ΔG°H2O invested in the shift in p*K*a exceeds the global stability of the protein.

The apparent p*Ka* measured for Lys-92 by linkage analysis corresponds to the free energy to transition between the fully folded, or closed state, and an alternative, open conformation in which the Lys can be hydrated when charged. In the case of Lys-92 this alternative, open conformation corresponds to the acid-unfolded state. The situation is different in variants with Lys-66. The NMR spectroscopy evidence from chemical shift perturbations at the site of Lys-66 and loss of signal (due to exchange broadening) in the surrounding regions strongly supports the interpretation that the protein undergoes equilibrium conformational changes into a state that retains a significant amount of structure while the C-terminus of helix-1 unfolds locally to expose the charged Lys-66 to water (Fig. 7 and supporting information). In all three cases with internal Lys-66 studied, the same local region of the protein was affected by the ionization of Lys-66. These NMR spectroscopy data support previous evidence from crystal structures and other studies with Lys-66 and Glu-66, that indicate that the local unwinding or over-winding of the C-terminus of helix-1 is all that is required for the ionizable moiety of Lys-66 to contact bulk water.^20,22,33,36^

All three proteins with Lys-66 populate the same locally unfolded conformations when Lys-66 is charged (Fig. 7 and supporting information). However, because the three proteins with Lys-66 differ in thermodynamic stability, the extent to which the native-like, locally unfolded conformation is populated in solution below the apparent p*Ka* value of Lys-66 differs for the three proteins. This follows the same trend as the differences in the stabilities of the closed states below the p*Ka* for the Lys, as determined from the linkage analysis (Table 1), and the NMR spectra of the amide backbone that were collected in the pH range below the apparent p*Ka* value (supporting information).

For the destabilized variant with Lys-66, the open state has similar ΔG as the reference protein with Lys-66 (difference of -0.5 kcal/mol). Although these variants populate a locally unfolded conformation in the range of pH of 6.3-7.0, the signal of the amide backbone is lost in a manner consistent with acid unfolding as the pH continues to decrease, beginning at pH 5.9 (supporting information). For the stabilized variant with Lys-66, both the open and closed states were stabilized by 1.9 kcal/mol relative to the reference protein, and there was no additional loss of signal from the protein backbone observed at pH values of 5.0-5.7, at or below the p*Ka* value. The fact that the same open conformation is populated is an important detail to consider given that the p*Ka* of Lys-66 was shifted to 7.0 (for the destabilized variant) relative to the p*Ka* of 5.7 in the reference protein), but the p*Ka* was unaffected in the stabilized variant). The absence of a shift in the p*Ka* of Lys-66 in the stabilized protein relative to the p*Ka* in the reference background protein is interesting. As shown in Fig. 9, this suggests that the difference in stability between the closed and open states was the same in the reference and in the stabilized forms of SNase with Lys-66. This is consistent with a situation in which mutations that alter the global stability of the protein in the native conformation affect the stability of every partially unfolded microstate in the ensemble that is composed of the same subset of native contacts ^56,57^

**Figure 9.**
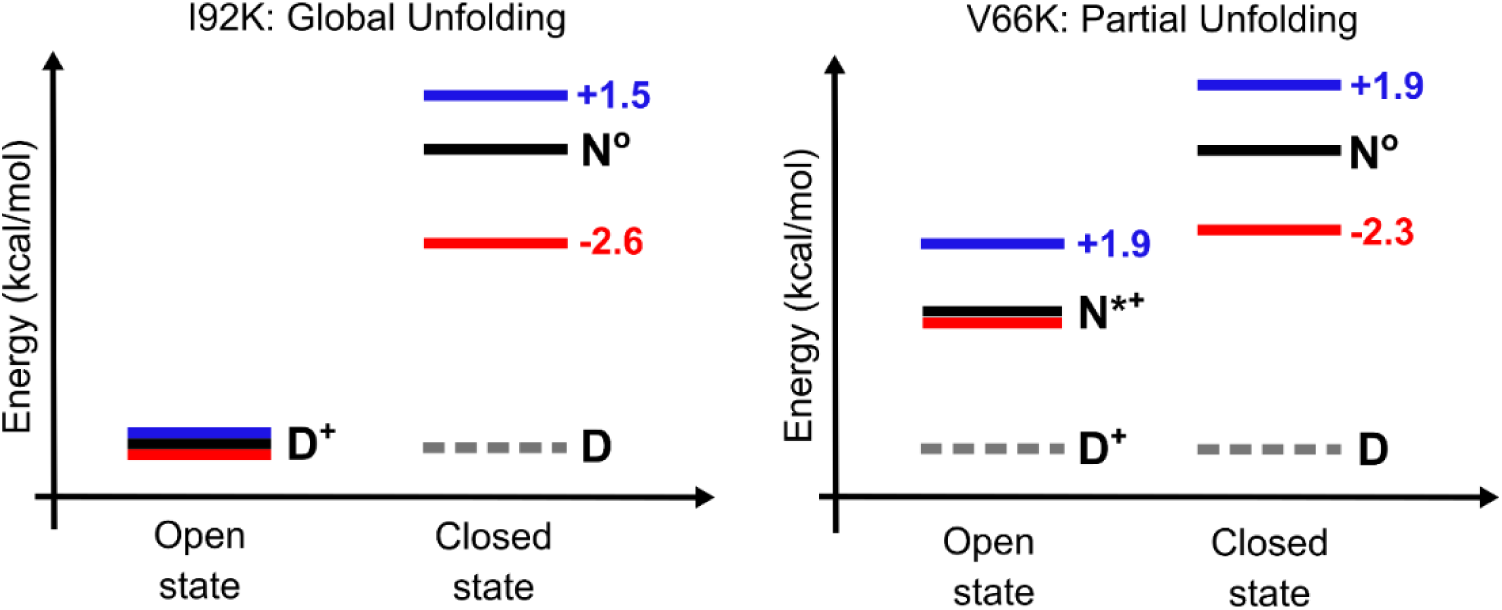
Free energy diagrams of the closed and open states in solution for proteins with stabilizing or destabilizing mutations, with Lys-92 **(left)** or with Lys-66 **(right)**. The stabilized Lys variants (blue), the destabilized Lys variants (red) are shown. The numbers refer to free energy changes calculated for the closed and open states with respect to the reference protein, shown in black. The closed state is calculated with the equation: ΔG^°^ ^𝐶𝐿𝑂𝑆𝐸𝐷^ = ΔG^°^ ^𝐵𝐺^ −ΔΔG_𝑚𝑢𝑡_, and the open state was calculated with the equation: ΔG^°^ ^𝑂𝑃𝐸𝑁^ = ΔG^°^ ^𝐶𝐿𝑂𝑆𝐸𝐷^ − 2.303 𝑅 𝑇(𝑝𝐾_𝑎,𝑤𝑎𝑡𝑒𝑟_ − 𝑝𝐾_𝑎_). In the closed state (at solution pH above p*Ka,w*), all Lys are neutral, and the fully folded, native conformation with the buried Lys (N°) is favorably populated over the denatured conformation (D). In the open state (at solution pH below the p*Ka*), the Lys-92 proteins transitioned to globally unfolded, denatured-like conformations (D^+^), and the Lys-66 proteins transitioned to partially unfolded, native-like conformations (N*^+^) that allow for the charged form of Lys to be hydrated.

The observations reported in this paper have important implications for structure-based calculations of electrostatic free energy in proteins. The properties of buried Lys, Glu, and Asp observed in SNase^13–15^ are representative of the properties of buried ionizable groups in other proteins.^58–61^ In SNase, the p*Ka* values of buried ionizable residues are governed by the propensity of the protein to undergo conformational reorganization coupled to the ionization of the buried group. The reorganization events are themselves determined by the global thermodynamic properties of the protein^39,62,63^ because that is what determines the probability that an alternative conformation will be populated (relative to all other states). This is fully consistent with NMR spectroscopy data^22,25–27,30,31,49^ and even crystal structures^18,20,21,27–29,32,33,64^ showing that in the cases described in this study, and in several others, the measured p*Ka* values are apparent p*Ka* values that are governed by pH-driven conformational transitions coupled to the ionization of the buried ionizable group.

These observations suggest that it will be very difficult to capture p*Ka* values of these and other internal ionizable groups with continuum electrostatic methods based on static crystal structures because the essential conformational reorganizational events that govern these p*Ka* values are very difficult to treat correctly in simulations with static structures.^65–67^ Corrections to the static nature of the starting structure with Monte Carlo sampling are unlikely to improve the calculations because sampling of rearrangements of the backbone with these methods remains extremely challenging.^53,68,69^ Methods that use molecular dynamics to consider relaxation of the protein backbone and side chains are possible, but these too suffer from shortcomings.^70–72^

Our observations with Lys-66 and Lys-92 in SNase suggest that accurate calculation of p*K*a values of internal ionizable groups in proteins will require methods that can both identify specific alternative conformations (local, sub-global, global) in the protein ensemble and predict free energy differences between conformational states with exquisite accuracy comparable to the energy reflected in the shifts of p*K*a values. This remains a considerable challenge for even the most sophisticated methods for structure-based energy calculations. For this reason, accurate treatment of the coupling between ionization of buried residues and conformational reorganization remains a challenge.^73,74^

These studies underscore how the hydration of ionizable groups acts as an important organizing force in macromolecular systems. Because the hydration energy of a single ionizable group is greater than the stability of most biological macromolecules or intermolecular interfaces,^75^ a single charge placed in a hydrophobic environment can be sufficient to modulate macromolecular systems and to drive transitions between conformational or assembly states. Biological function of proteins always involves transitions between states. In the case of the most fundamental energy transduction processes in biological systems in which proteins manage high energies, the essential recurring structural motif consists of an ionizable residue(s) with an anomalous p*K*a buried in a hydrophobic, dehydrated environment, poised to react to a change in pH with a coupled equilibrium between H^+^ binding/release and conformational transitions. Lys-62 and Lys-99 in SNase are excellent mimics of the fundamental energy transduction motif in biological systems.^76,77^

## ACKNOWLEDGEMENTS

NMR spectroscopy experiments were performed in the BioNMR facility at Johns Hopkins University. The authors with to thank Profs. Doug Barrick, Juliette Lecomte, Vince Hilser, and Dominique Frueh, for their helpful suggestions.

