## Supporting Information for "The p*K_a_* values of buried ionizable groups in proteins can be determined by the thermodynamic stability of the protein"

**Supporting Information:** The p*K_a_* values of buried ionizable amino acids can be determined by the thermodynamic stability of the protein

**Supporting Figures**

1. ^1^H^15^N-HSQC spectrum at pH 7 of the stabilized background protein overlaid with the spectrum of the reference protein
2. Combined HN CSPs calculated between pH 5 and 7 for the stabilized and reference background proteins
3. ^1^H^15^N-HSQC spectrum at pH 7 of the destabilized background protein overlaid with the spectrum of the reference protein
4. ^1^H^15^N-HSQC spectrum of the stabilized protein with Lys-92 at pH 5.3 overlaid with the spectrum collected at pH 4.3
5. ^1^H^15^N-HSQC spectra of the pH titration of the stabilized background protein
6. ^1^H^15^N-HSQC spectra of the pH titration of the stabilized background protein with Lys-66
7. ^1^H^15^N-HSQC spectra of the pH titration of the destabilized background protein
8. ^1^H^15^N-HSQC spectra of the pH titration of the destabilized background protein with Lys-66
9. Calculation of combined HN ∆CSPs for the stabilized protein with Lys-66
10. Calculation of combined HN ∆CSPs for the destabilized protein with Lys-66
11. 2D (HA)CACO spectra of the pH titration of the stabilized background protein with and without Lys-66
12. Calculation of the combined CaCO ∆CSPs for the stabilized protein with Lys-66
13. Comparison of the HN and CaCO (∆)CSPs determined for stabilized and destabilized proteins with and without Lys-66 as mapped onto the structure of the reference protein
14. Residues identified with a pH-dependent loss of signal in ^1^H^15^N-HSQC spectra for the destabilized background protein with and without Lys-66 mapped onto the structure of the reference protein with or without Lys-66

**Supplemental Tables**

1. Thermodynamic parameters of the substitutions made at the protein surface to engineer the (de)stabilized background proteins from the reference protein
2. Residues that were identified from calculation of (∆)CSPs from NMR spectra for the (de)stabilized proteins with and without Lys-66
3. Residues identified to lose signal at different pH values below the p*K_a_* value of 7.0 for Lys-66 in the destabilized protein as determined from NMR spectra and as mapped onto the structures shown in Supp. Fig. 14

**Additional Material and Methods:**

1. Manual titration procedure used for determination of the stabilities of the background protein candidates shown in supplemental table 1

**FIGURES:**

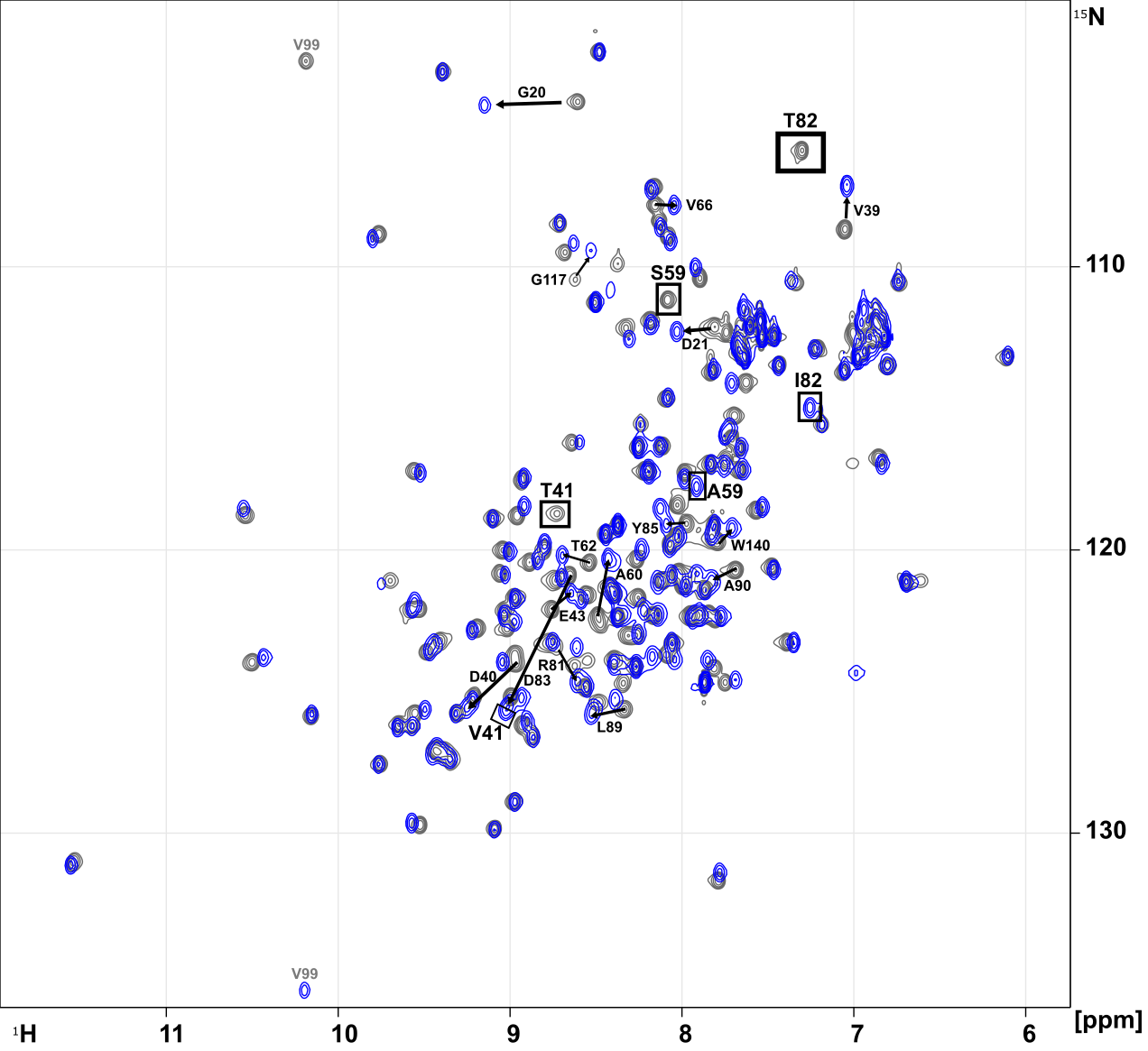

**Supp. Fig. 1.** ^1^H^15^N-HSQC spectrum of the stabilized background protein (∆+PHS T41V/S59A/T82I) overlaid with the spectrum of the reference protein (∆+PHS, as collected and published by Castañeda et al.^1^) at pH 7. V99 is labelled in gray to show how the peak folds into the two spectra at different positions. The peaks that pertain to the mutations that were made are shown as boxes. Labelled arrows indicate positions that exhibit altered chemical shifts because of the mutations made.

The residues that exhibit changes to the ^1^H,^15^N chemical shift (and thus the chemical microenvironment) are primarily i±3 of the substitutions made to engineer the background proteins. The change in CSPs between pH 7 and 5 was analyzed for the stabilized protein compared to the reference background protein (shown in Supp. Fig. 2), which validates that the effect that the mutations have on the structure of the protein are ineffectual.

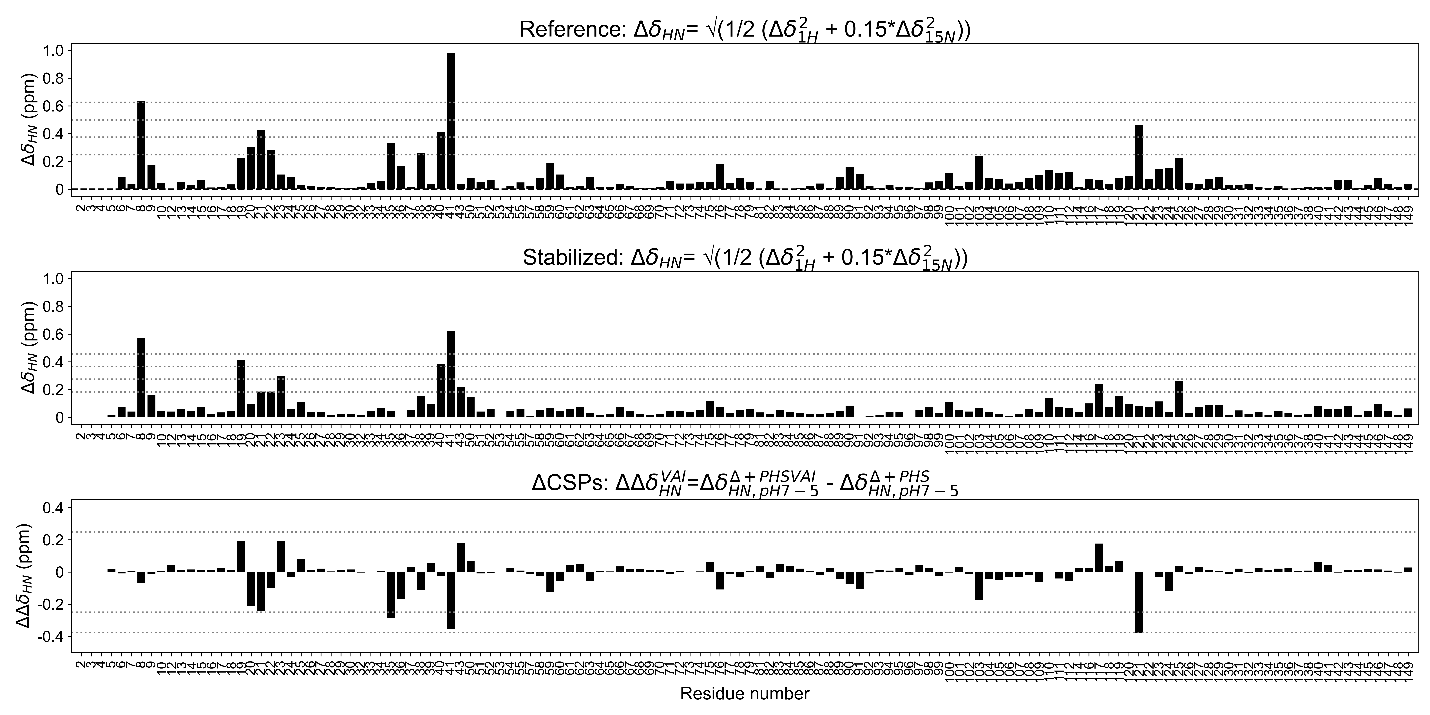

**Supp. Fig. 2.** The combined HN CSPs calculated per residue between pH 7 and 5 for the reference protein (∆+PHS, **TOP**) and stabilized background protein (∆+PHS VAI, **MIDDLE**). Dashed lines are 1-5 times the standard deviation on the mean with the propagation of error (1σ-5σ). **(Bottom)** Change in the combined ^1^H,^15^N CSPs between the stabilized and reference background proteins to evaluate if residues undergo a pH-dependent conformational change, relative to the reference protein, prior to the addition of a buried Lys. R35, T41, and H121 are the only positions that have a significant change in ∆CSPs, which are negative and suggest that the pH-dependent response is suppressed relative to the reference protein. In whole, this is consistent with an increase in stability of the protein.

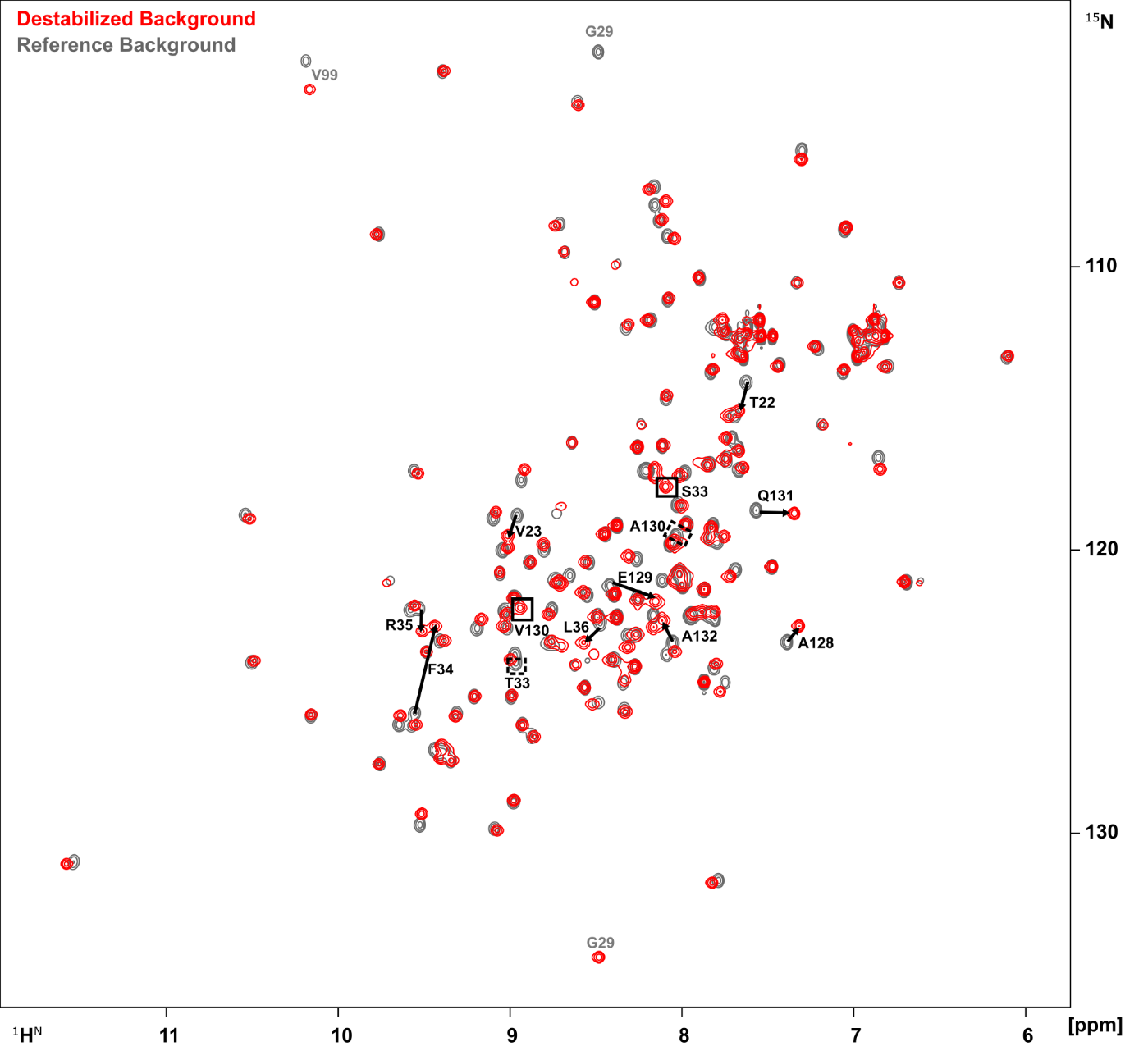

**Supp. Fig. 3.** ^1^H^15^N-HSQC spectrum at pH 7 of the destabilized background protein (∆+PHS

T33S/A130V) overlaid with the spectrum of the reference protein (∆+PHS, as collected and published by Castañeda et al.^1^). G29 and V99 are labelled in gray because these peaks fold into the two spectra at different positions. The peaks that pertain to the mutations that were made are shown as boxes. Labelled arrows indicate positions that exhibit altered chemical shifts.

The residues that exhibit changes to the ^1^H,^15^N chemical shift are i±3 of the substitutions made to engineer the background proteins, except T22 and V23 that interacts with T33S on the adjacent beta strand in the beta barrel of SNase. This indicates that the effect that the mutations have on the structure of the protein are local and at the surface.

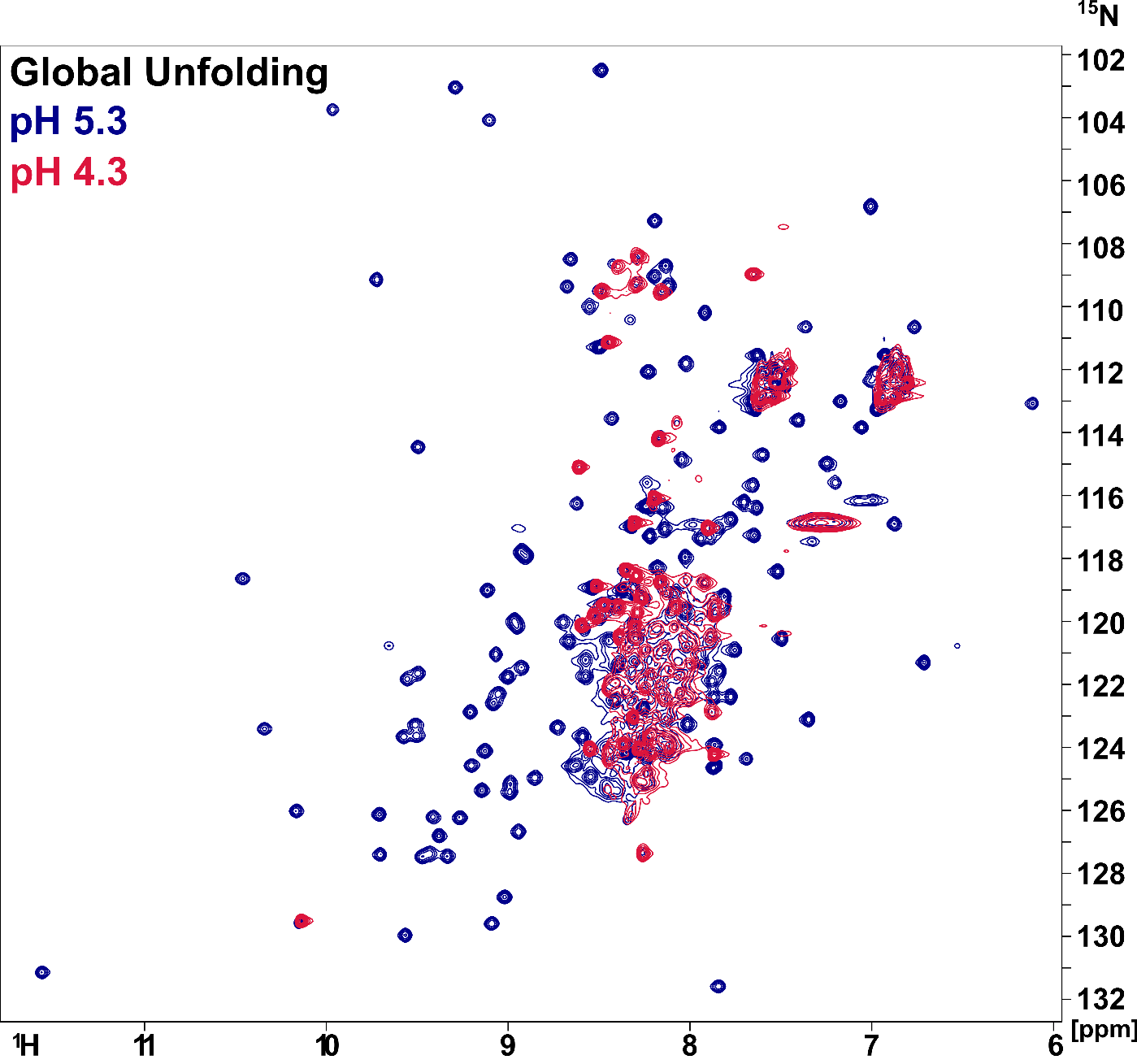

**Supp. Fig. 4.** An overlay of the ^1^H^15^N-HSQC spectrum of the stabilized protein with Lys 92 at pH 5.3 in dark blue, where the protein is largely folded but enters a slow exchange regime with a globally unfolded population, and at pH 4.3 in red, where the protein has transitioned completely to a globally unfolded state. The pH of 5.3 is within error of the p*K_a_* value of Lys-92 in the reference protein, and pH 4.3 is below the p*K_a_* value of Lys-92 measured in the stabilized protein.

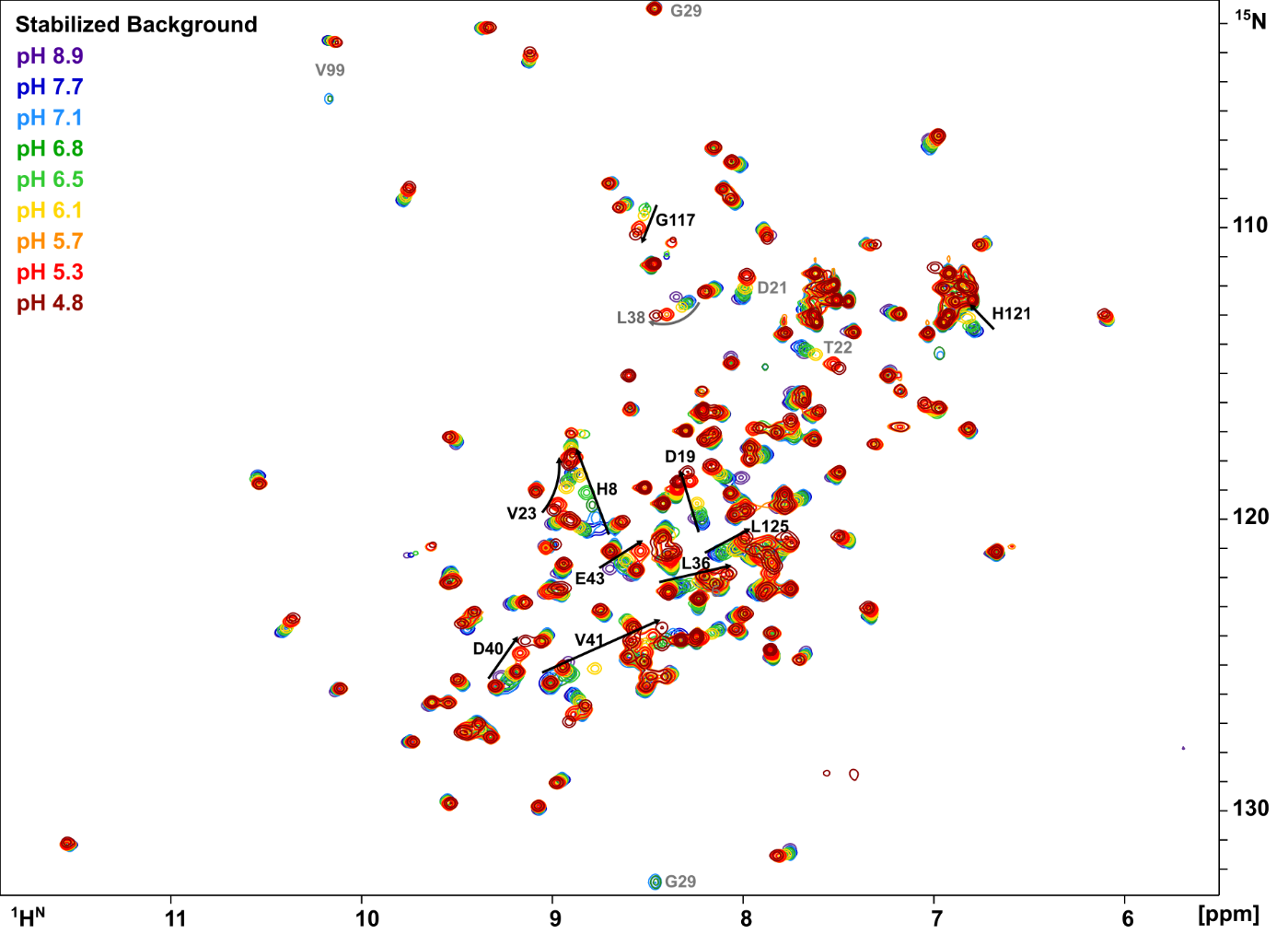

**Supp. Fig. 5.** ^1^H^15^N-HSQC spectra of the pH titration of the stabilized background protein. Arrows indicate residues that exhibit chemical shift perturbations (CSPs) between pH 7.1 and 4.8. Residues that are labeled gray appear to titrate in this pH range, but CSPs are insignificant. Residues G29 and V99 are labeled gray to denote how these peaks are folded differently into the spectra.

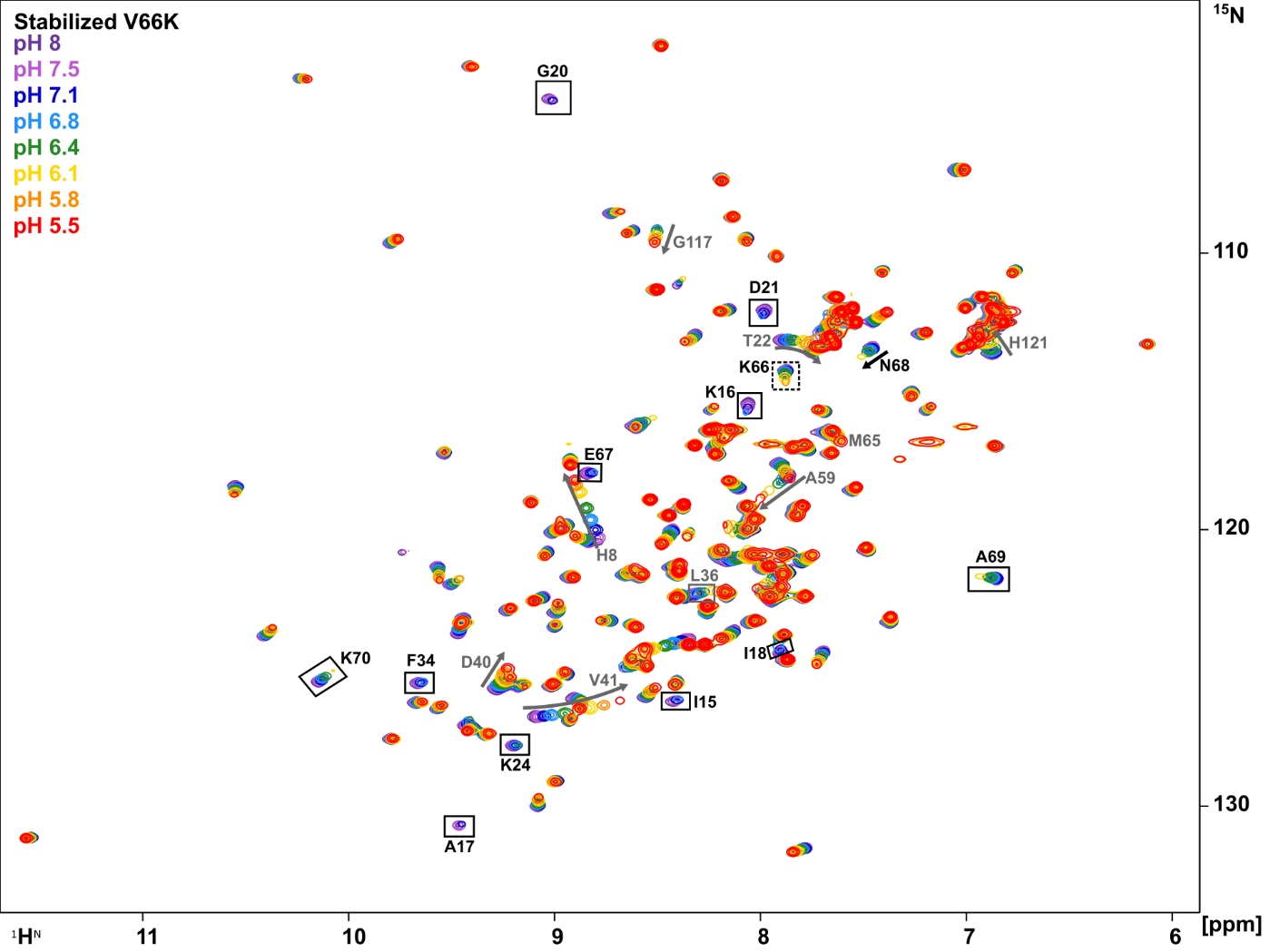

**Supp. Fig. 6.** ^1^H^15^N-HSQC spectra of the pH titration of the stabilized protein with Lys 66. Lys-66 is indicated by a dashed box. Residues that underwent a loss of signal below pH 6.1-7.1 are labelled with solid boxes. The residues that underwent CSPs are shown labelled with arrows. The residues are labeled that are affected by the ionization of Lys-66 at its p*K_a_* value of 5.7 (black font) or insignificant after consideration of the CSPs of the background protein over the same pH range (gray font).

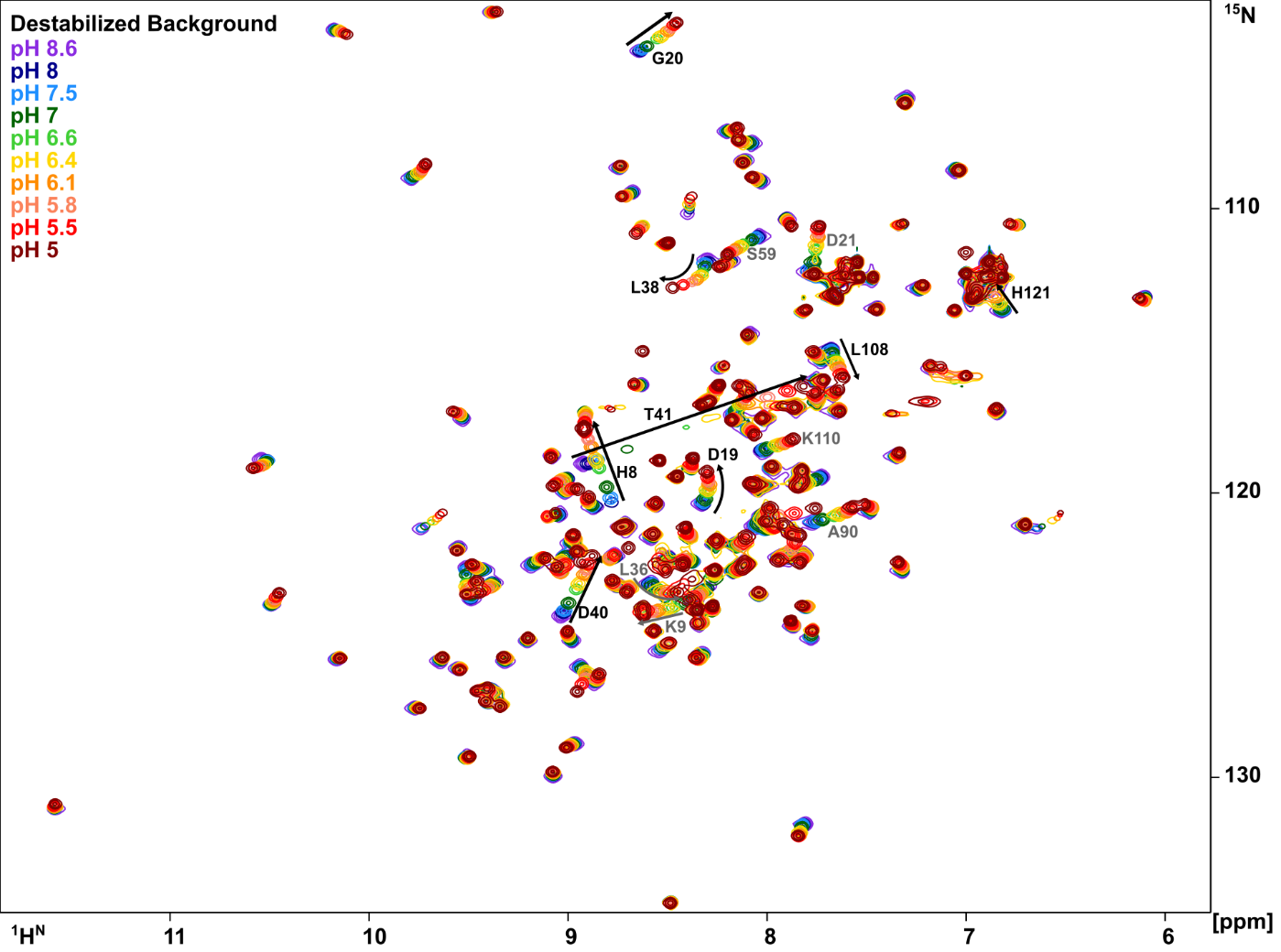

**Supp. Fig. 7.** ^1^H^15^N-HSQC spectra of the pH titration of the destabilized background protein. Arrows indicate residues that exhibit significant CSPs (>2σ) between pH 8 and 6.3. Residues that are labeled gray titrate in this pH range, but CSPs are insignificant.

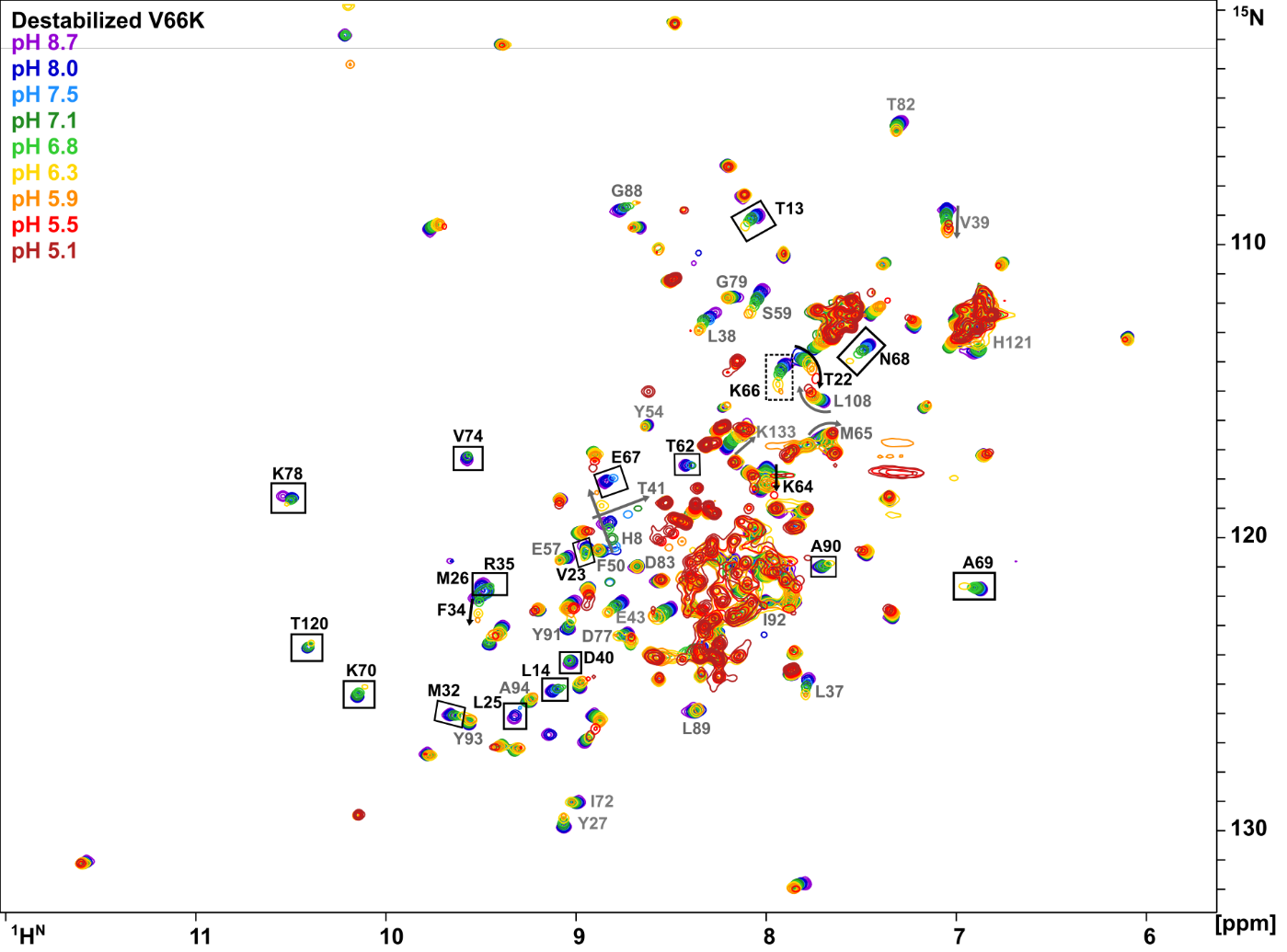

**Supp. Fig. 8.** ^1^H^15^N-HSQC spectra of the pH titration of the destabilized protein with Lys 66. Lys-66 is indicated by a dashed box. The residues are labeled that are affected by the ionization of Lys-66 at its p*K_a_* value of 7.0 (black font). Residues that lose signal below pH 6.3 are labelled with solid boxes, and those in gray font lose signal below pH 5.9. The residues that exhibit CSPs are shown with arrows, where those in black font are significant and those in gray font are insignificant after the consideration of the background protein.

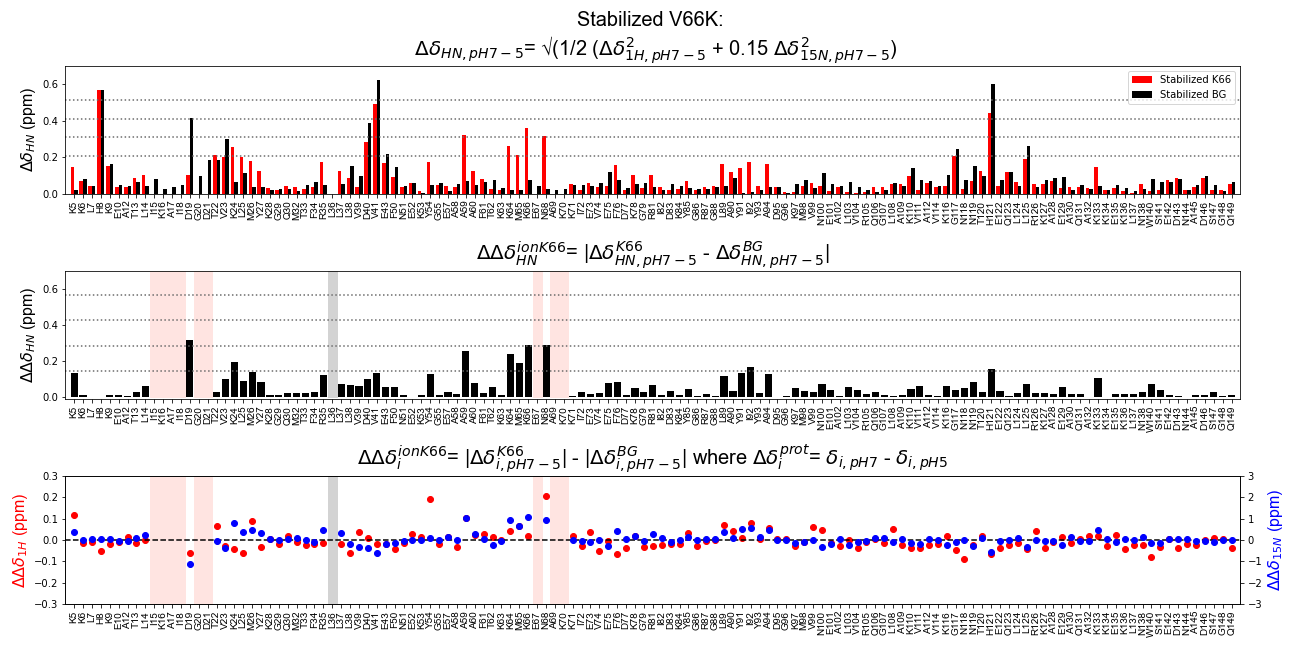

**Supp. Fig. 9.** Calculation of the pH-dependent change in the chemical shift perturbations (∆CSPs) of ^1^H,^15^N nuclei for the stabilized protein with Lys-66. Only residues that were assigned in one of the two pH spectra are labeled on the x-axis, where lack of data denotes residues that lost signal at either pH value. The vertical bands designate residues that lost signal with the change in pH in either the protein with Lys-66 (light red) or the stabilized background protein (gray). **(TOP)** Combined HN CSPs calculated per residue between pH 7.1 and 5.0 for the stabilized protein with Lys-66 (red) and between pH 7.1 and 4.8 for the stabilized background protein (black). Dashed lines are 2-5 times the standard deviation on the mean. **(MIDDLE)** The change in the combined HN CSPs that were determined from subtracting the background protein from the Lys-66 variant. Dashed horizontal lines are the propagated error of the standard deviation (1σ-5σ). **(BOTTOM)** Change in ^1^H (red, left axis) and ^15^N (blue, right axis) CSPs calculated separately to demonstrate if one or both nuclei contribute to the combined CSP calculation above.

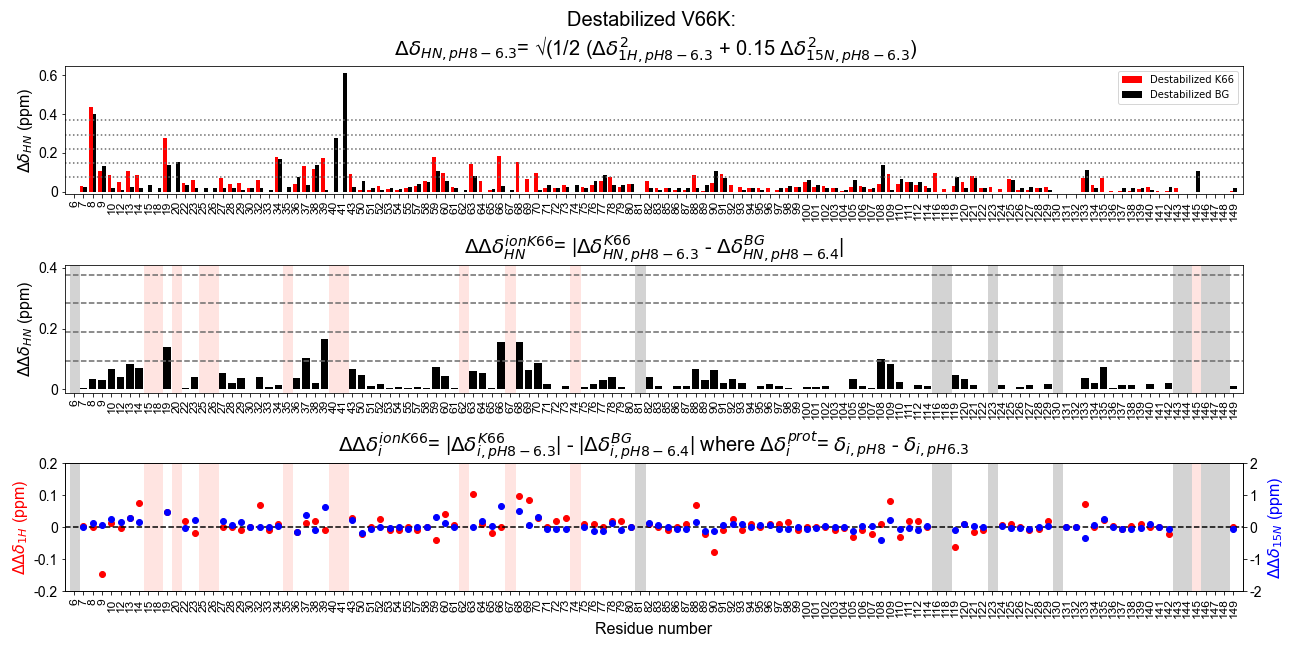

**Supp. Fig. 10.** Calculation of the pH-dependent change in the chemical shift perturbations (∆CSPs) for ^1^H^15^N nuclei for the destabilized protein with Lys-66. For the three plots, only residues that were assigned in one of the two pH spectra are labeled on the x-axis, where lack of data denotes residues that lost signal. The vertical bands designate residues that lost resonance in either the destabilized protein with Lys-66 (light red) or the destabilized background protein (gray). **(TOP)** Combined HN CSPs calculated per residue between pH 8.0 and 6.3 for the Destabilized Lys-66 protein (red) and destabilized background protein (black). Dashed lines are 1-5 times the standard deviation on the mean. **(MIDDLE)** Change in combined HN CSPs subtracting the background protein from the Lys-66 variant. Dashed lines are the standard deviations with the propagation of error (1σ-5σ). **(BOTTOM)** Change in ^1^H (red, left axis) and ^15^N (blue, right axis) CSPs calculated separately to demonstrate if one or both nuclei contribute to the combined CSP calculation.

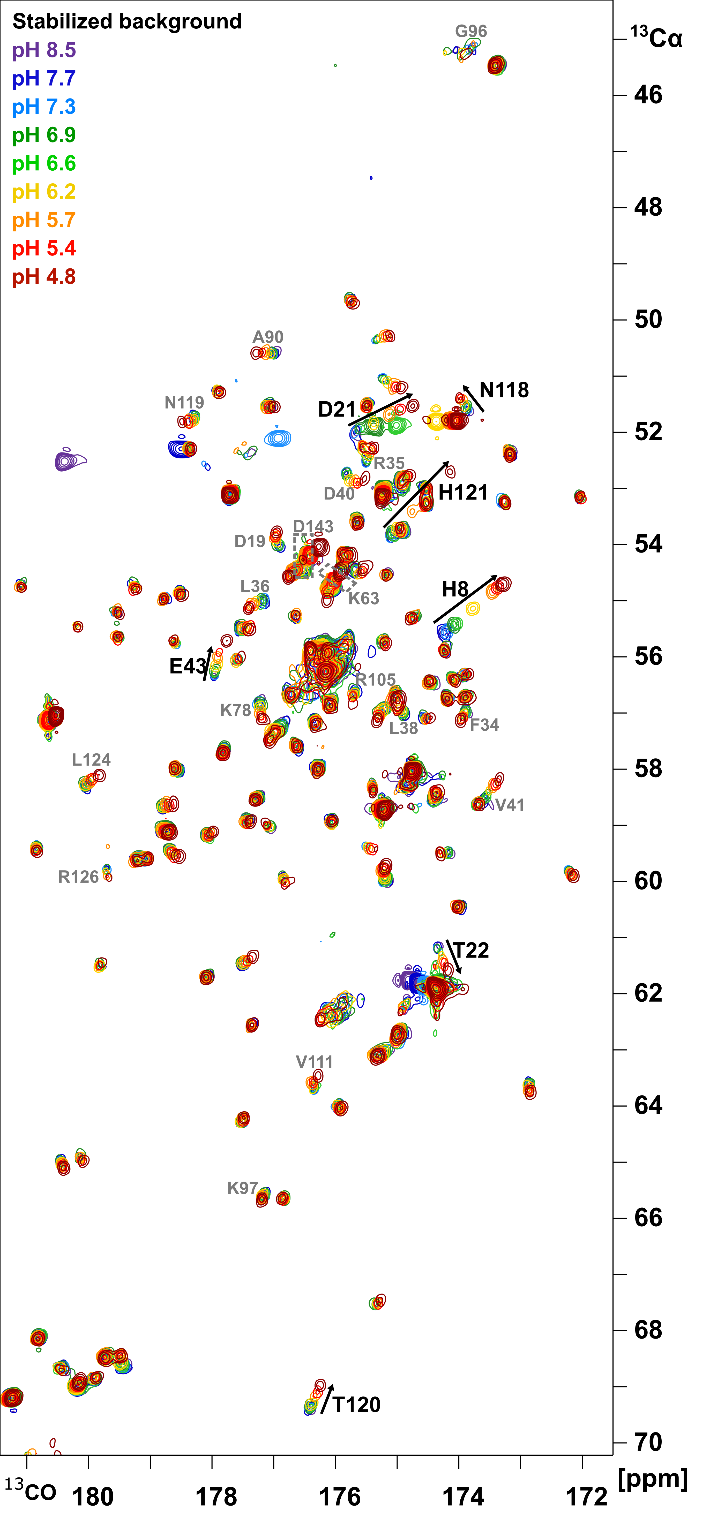

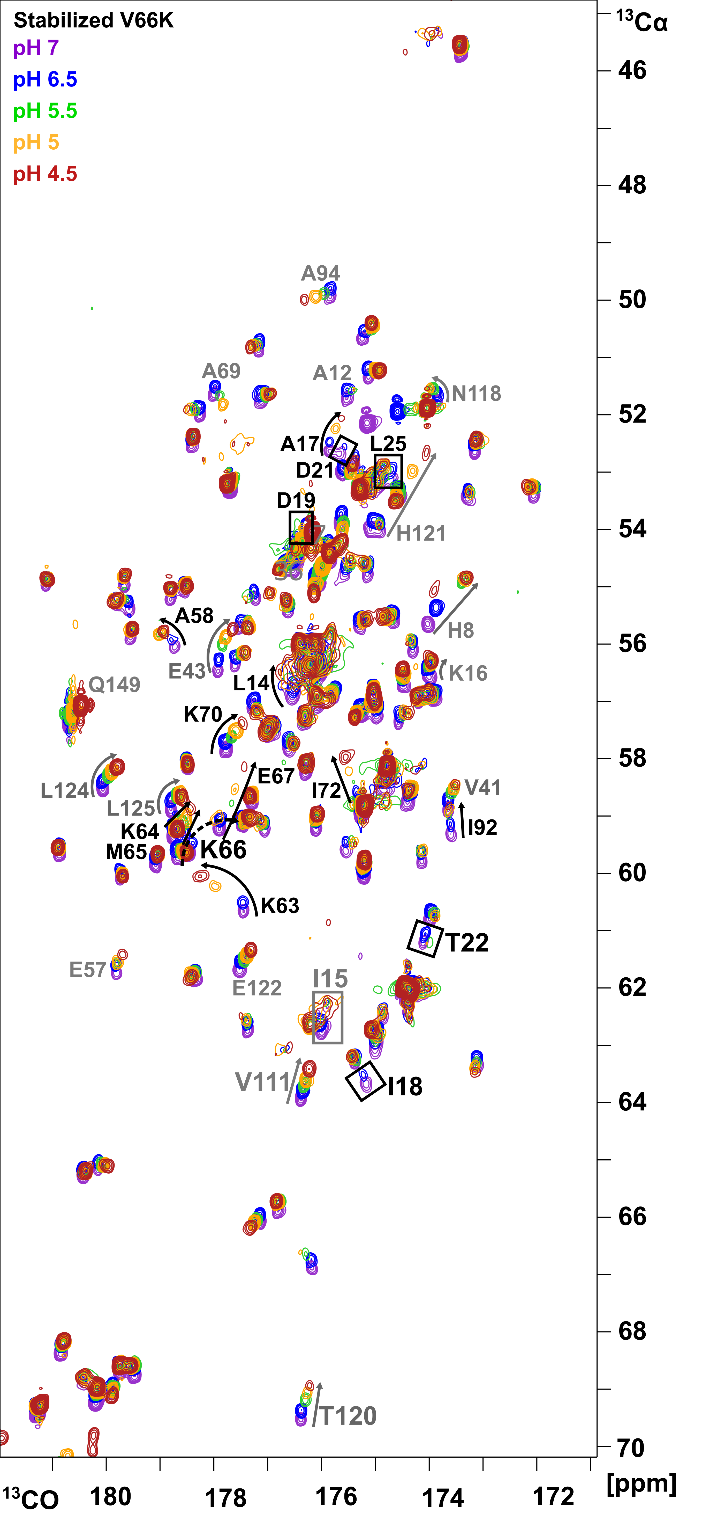
**Supp. Fig. 11.** 2D ^13^C-detect (Hα)CαCO spectra collected as a pH titration for the stabilized background protein (left) and the stabilized protein with Lys-66 (right). **(LEFT)** Arrows indicate residues that exhibit CSPs between pH 6.9 and 4.8. Residues that are labeled gray titrate in this pH range, but CSPs are insignificant. **(RIGHT)** Lys-66 is indicated by a dashed arrow. The residues that are labeled in black font are the residues that are affected by the ionization of Lys-66 at its p*K_a_* value of 5.7. Residues that lose signal are labelled with boxes. The residues that have CSPs are shown labelled with arrows, where those in gray font are insignificant after subtraction of the background protein.

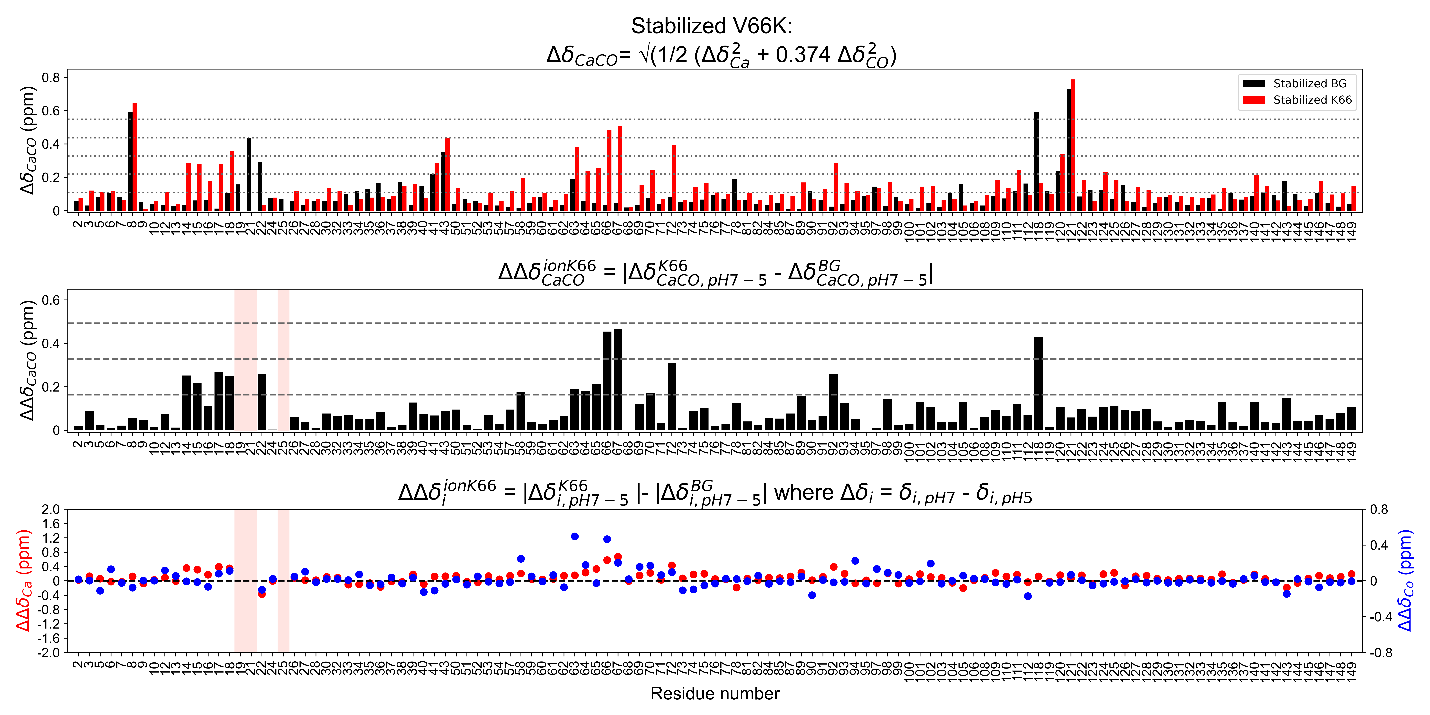

**Supp. Fig. 12.** The combined Cα,CO chemical shift perturbations (∆CSPs) for the stabilized protein with Lys-66. For the three plots, only residues that were assigned in one of the two pH spectra are shown on the x axis, such that lack of data denotes residues that lost signal, also noted by light red bars. **(TOP)** Combined Cα,CO CSPs calculated per residue between pH 7.0 and 5.0 for the stabilized protein with Lys-66 (red) and the stabilized background protein (black). Dashed lines are 1-5 times the standard deviation on the mean. **(MIDDLE)** The change in the combined Cα,CO CSPs that were calculated by subtracting the background protein from the Lys-66 variant. Dashed lines are 1-3 times the standard deviation with the propagation of error. **(BOTTOM)** The change in ^13^Cα (red, left y-axis) and ^13^CO (blue, right y-axis) pH-dependent CSPs calculated separately.

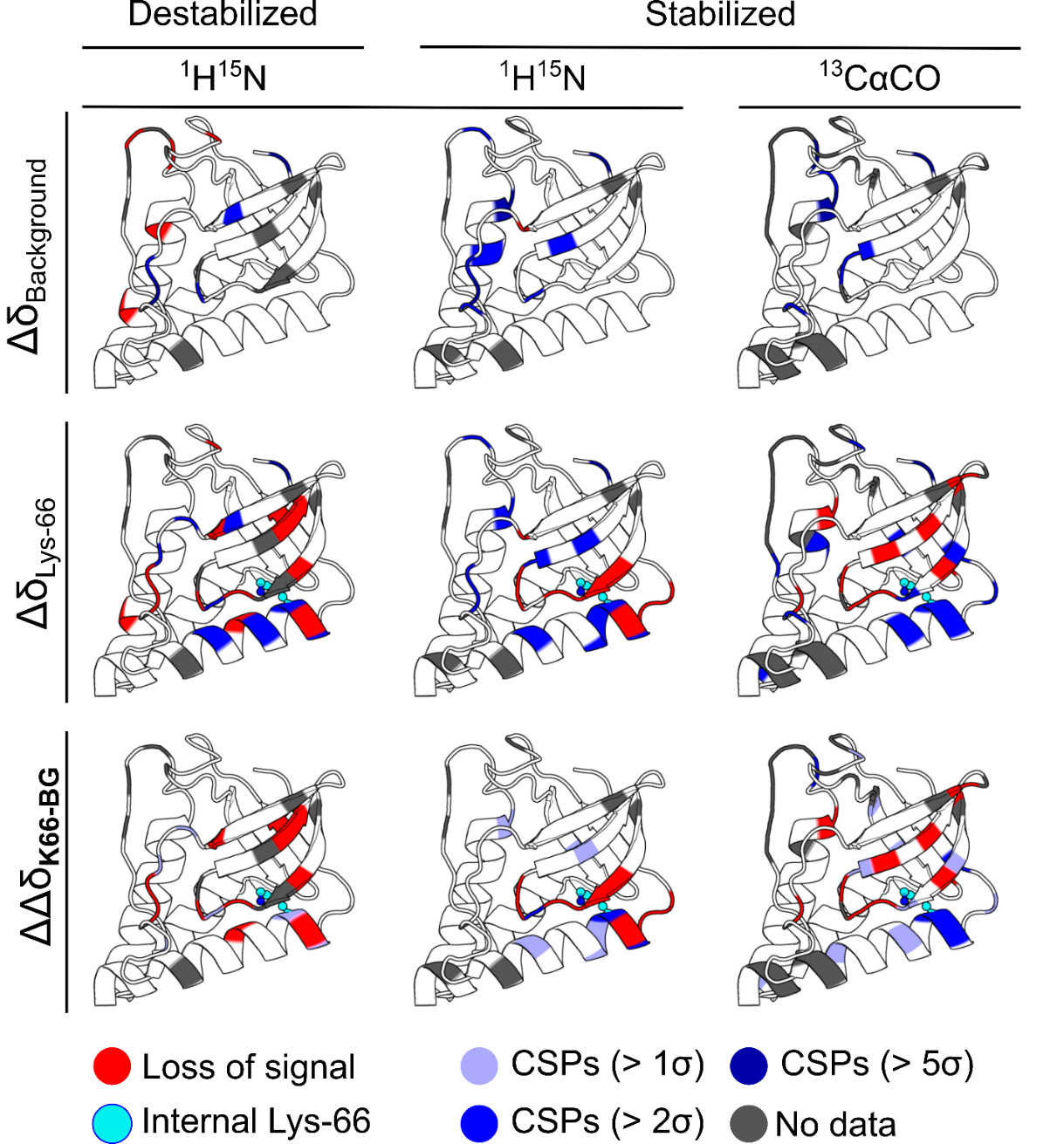

**Supp. Fig. 13.** Comparison of the ^1^H^15^N (∆)CSPs calculated for the destabilized and stabilized proteins with and without Lys-66 to the CαCO CSPs of the stabilized protein with and without Lys-66. Residues that underwent loss of signal at 0.7 pH units below the p*K_a_* value of Lys-66 are shown in red, and residues that experience CSPs (greater than 1σ-5σ) are shown in shades of blue. Residues that lack assignments are indicated in gray. The side chain of internal Lys-66 is shown in cyan.

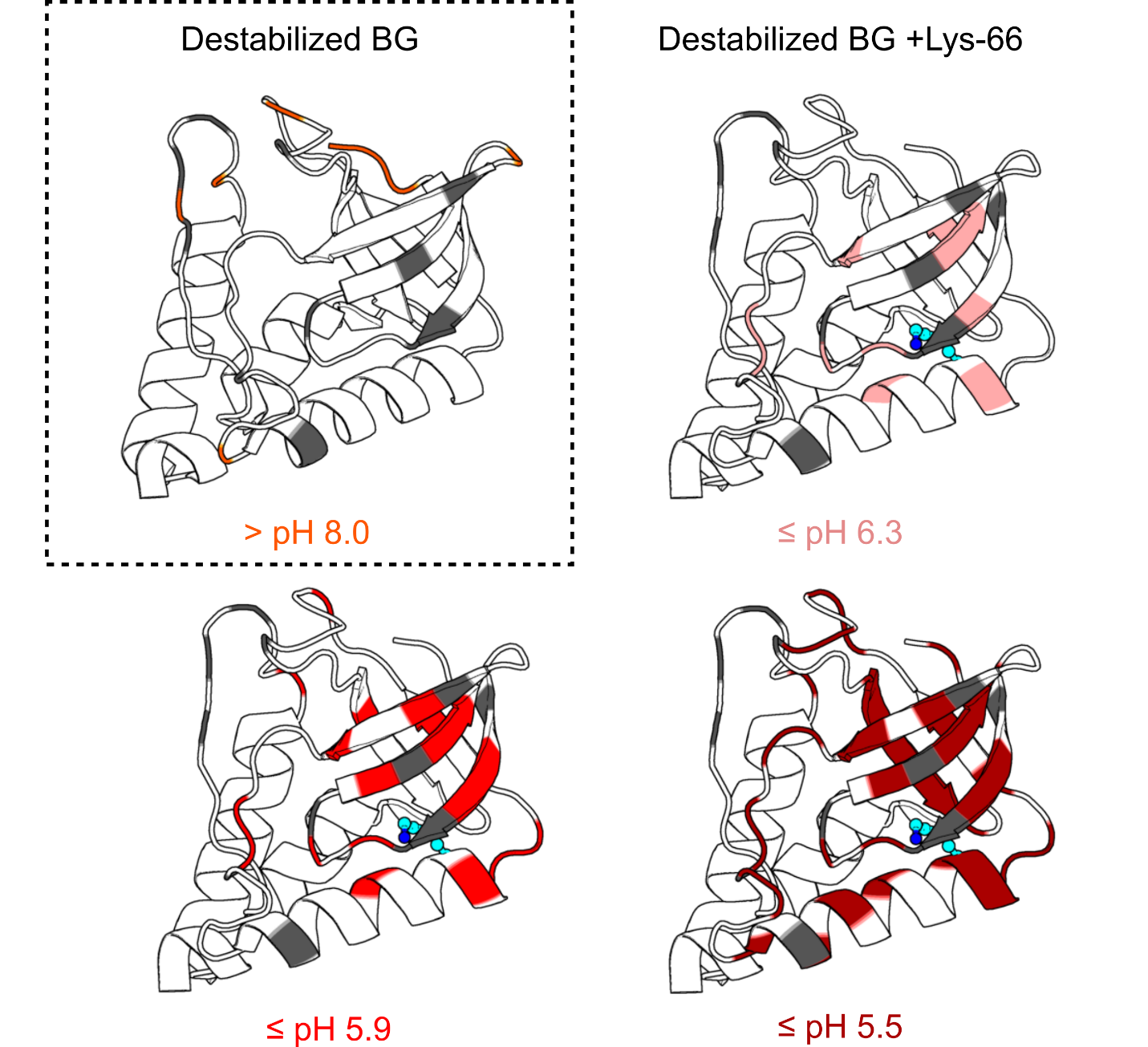

**Supp. Fig. 14.** For the destabilized protein with and without Lys-66, residues that undergo a pH-dependent loss of signal in ^1^H^15^N-HSQC are shown mapped onto the crystal structures of the reference protein with or without Lys-66. Loss of signal occurred either above pH 8 due to base-catalyzed exchange as shown for the destabilized protein without Lys-66 (orange) or below the p*K_a_* value of 7.0 for the destabilized protein with Lys-66 (light red: < pH 6.3, red: < pH 5.9, dark red: < pH 5.5; see Supp. Fig. 7-8 and Supp. Table 3). Residues that lack assignments are indicated in gray. The side chain of internal Lys-66 is shown in cyan, and the nitrogen moiety is blue.

**SUPPLEMENTAL TABLES:**

**Supp. Table 1.** The thermodynamic parameters of the substitutions made at the protein surface to engineer the (de)stabilized background proteins from the reference protein

| Protein variants | | ∆G°_H2O_^a^  kcal mol^-1^ | ∆∆G_var-ref_ ^b^  kcal mol^-1^ | m-value^c^  kcal  mol^-1^ M^-1^ | $\frac{m_{var}}{m_{ref}}$^d^ | ∆G_coupling_^e^  kcal mol^-1^ |
| --- | --- | --- | --- | --- | --- | --- |
| Reference | ∆+PHS | -11.9 ± 0.1 | 0 | 4.9 ± 0.1 | 1.00 | 0 |
| Stabilized | ∆+PHS T41V | -12.3 ± 0.2 | 0.4 ± 0.3 | 4.7 ± 0.1 | 0.96 | 0.4 |
|  | ∆+PHS S59A | -12.6 ± 0.2 | 0.7 ± 0.3 | 4.7 ± 0.1 | 0.96 | 0.7 |
|  | ∆+PHS T82I | -11.9 ± 0.1 | 0.0 ± 0.2 | 4.7 ± 0.1 | 0.96 | 0.0 |
|  | ∆+PHS T41V,S59A | -13.7 ± 0.2 | 1.8 ± 0.2 | 4.7 ± 0.1 | 0.96 | 0.7 |
|  | ∆+PHS T41V,T82I | -12.8 ± 0.1 | 0.9 ± 0.2 | 4.8 ± 0.1 | 0.98 | 0.5 |
|  | ∆+PHS S59A,T82I | -12.7 ± 0.1 | 0.8 ± 0.2 | 4.7 ± 0.1 | 0.96 | 0.1 |
|  | ∆+PHS T41V,S59A,T82I | -13.1 ± 0.1 | 1.3 ± 0.2 | 4.6 ± 0.1 | 0.94 | -1.2 |
| Destabilized | ∆+PHS T33S | -10.4 ± 0.1 | -1.5 ± 0.2 | 4.9 ± 0.1 | 1.00 | -1.5 |
|  | ∆+PHS A130V | -10.8 ± 0.1 | -1.1 ± 0.2 | 4.8 ± 0.1 | 0.98 | -1.1 |
|  | ∆+PHS T33S,A130V | -9.8 ± 0.1 | -2.1 ± 0.2 | 5.1 ± 0.1 | 1.04 | 0.5 |

^a^The stabilities and m-values^c^ in water were measured at pH 7 from chemical denaturation experiments in Gdm in the presence of 100 mM KCl and 25 mM Tris buffer at 25°C.

^b^The difference in stability was calculated between the measured stability at pH 7 for the background protein with additional mutations (variant) and the pH-independent stability of the reference protein at pH 7. The error reported is the propagated experimental error.

^d^The ratio of the m-value of the variant relative to the reference protein. An experimental error in m of ± 0.1 kcal mol^-1^ M^-1^ is a 2% change in the m-value relative to the reference protein.

^e^The non-additive coupling energies were calculated for single substitution variants as: ∆G_coup,mut1_ = ∆G_REF_ - ∆G_VAR_, for double substitution variants as: ∆G_coup,mut12_ = ∆G_REF_ - ∆G_VAR,mut12_ - ∆G_coup,mut1_ - ∆G_coup,mut2_, and for the triple substitution variant as: ∆G_coup,mut123_ = ∆G_REF_ - ∆G_VAR,mut123_ - ∆G_coup,mut12_ - ∆G_coup,mut13_ - ∆G_coup,mut23_ - ∆G_coup,mut1_ - ∆G_coup,mut2_ - ∆G_coup,mut3_

**Supp. Table 2.** The residues identified from NMR spectra for the (de)stabilized proteins with and without Lys-66 as mapped onto the structures shown in Fig. 7 of the main text and Supp. Figures

| (∆)∆δ_nuclei,pH_  Protein variant | Loss of signal | Chemical Shift Perturbations | | | No data  (% assigned) |
| --- | --- | --- | --- | --- | --- |
|  |  | 1σ-2σ | > 2σ-5σ | > 5σ |  |
| ∆δ_HN,pH 7-5_  Stabilized BG +V66K | I15, K16, A17, I18, G20, D21, L36, E67, A69, K70 | K5, K9, V23, L25, M26, Y7, R35, L37, E43, Y54, A60, F76, R81, L89, A90, Y91, I92, A94, T120, Q123, L125, K133 | T22, K24, D40, V41, A59, K64, M65, K66, N68, G117, H121 | H8 | A1-T4, P11, P31, P42, P56, Q80, Y113, Y115, I139  (131 of 139 assignable residues, 94% assigned) |
| ∆δ_HN,pH 7-5_  Stabilized BG | L36 | K9, D21, T22, L25, L38, F50, E75, N100, K110, N119, Q123 | D19, V23, D40, E43, G117, L125 | H8, V41, H121 |  |
| ∆∆δ_HN,pH 7-5_  Stabilized BG +V66K | I15, K16, A17, I18, G20, D21, E67, A69, K70 | K24, A59, K64, M65, I92, H121 | D19, K66, N68 | - |  |
| ∆δ_HN,pH8-6.3_  Destabilized BG +V66K | K6, I15, I18, G20, L25, M26, R35, D40, T41, T62, E67, V74, R81, V130, N144, A145, S147, G148 | K9, E10, T13, L14, V23, Y27, M32, L38, E43, A60, A69, K70, K78, G88, Y91, A109, K116, H121, L125, K133, E135 | D19, F34, L37, V39, S59, K63, K66, N68 | H8 | A1-K5, P11, K16, A17, D21, K24, P31, P42, P56, K84, Y113, Y115, G117  (126 of 139 assignable residues, 90.6% assigned) |
| ∆δ_HN,pH8-6.3_  Destabilized BG | K6, R81, K116, N118, Q123, V130, D143, N144, D146, S147, G148 | K9, D19, L36, L38, S59, K63, D77, A90, I108, N119, K133, A145 | G20, F34, D40 | H8, T41 |  |
| ∆∆δ_HN,pH8-6.3_  Destabilized BG +V66K | I15, I18, G20, L25, M26, R35, D40, T41, T62, E67, V74, A145 | D19, L37, V39, K66, N68, L108 | - | - |  |
| (∆)∆δ_nuclei,pH_  Protein variant | Loss of signal | CSPs  1σ-2σ | CSPs  > 2σ-5σ | CSPs  > 5σ | No data  (% assigned) |
| ∆δ_CαCO,pH 7-5_  Stabilized BG +V66K | D19, D21, L25 | S3, K6, A12, K16, M26, Q30, M32, L38, V39, F50, E57, A58, A69, V74, E75, L89, Y91, Y93, A94, K97, M98, E101, A102, A109, K110, N118, E122, L125, K127, A128, E135, W140, S141, D146, Q149 | L14, I15, A17, I18, E43, K63, K64, M65, K66, E67, K70, I72, I92, V111, T120, L124 | H8, H121 | A1-T4, P11, G20, V23, G29, P31, P42, G55, P56, G79, Q80, D83, G86, G88, G107, Y113, V114, Y115, K116, G117, N138, I139  (119 of 133 assignable residues, 89.5% assigned) |
| ∆δ_CαCO,pH 7-5_  Stabilized BG | - | D19, F34, R35, L36, L38, D40, V41, K63, K78, A90, K97, R105, V111, A112, N119, Q123, L124, R126, D143 | D21, T22, E43, T120 | H8, N118, H121 |  |
| ∆∆δ_CαCO,pH 7-5_  Stabilized BG +V66K | D19, D21, L25 | L14, I15, A17, I18, T22, A58, K63, K64, M65, K70, I72, I92 | K66, E67, N118 | - |  |

**Supp. Table 3.** The residues identified to lose resonance at different pH values below the p*K_a_* value of 7.0 for Lys-66 in the destabilized protein from NMR spectra, as mapped onto the structures shown in Supp. Fig. 14. Residues are bolded if they are new to the list at the lower pH.

| Destabilized BG +V66K | |
| --- | --- |
| ∆∆δ_1H15N,pH8-pH_ **_#_** | Loss of Signal |
| pH 6.3 | I15, I18, G20, L25, M26, R35, D40, T41, T62, E67, V74 |
| pH 5.9 | **T13, L14**, I15, I18, G20, **V23**, L25, M26, **M32**, R35, D40, T41, T62, E67, **A69, K70**, V74, **K78, A90, T120** |
| pH 5.5 | T13, L14, I15, I18, G20, V23, L25, M26, Y27, M32, **F34**, R35, **L37, L38**, D40, T41, **F50, E57, S59**, T62, E67, **N68,** A69, K70, **I72**, V74, **D77**, K78, **G79, T82, D83, G88, L89**, A90, **Y91, I92, Y93, A94**, T120 |

**Additional Materials and** **Methods:**

**Manual titrations:** Chemical denaturation experiments were performed at pH 7 as a manual titration for single, double and triple variants of the reference background protein that do not contain internal Lys residues (pertaining to the data in supplemental Table 1). The stabilities were additionally validated by measurement with an automated titration as described above. For manual titration experiments, buffer and titrant solutions were made to a volume of 60 mL that contained 100 mM KCl and 25 mM HEPES buffer without or with 6 M GdmCl, respectively. Protein was thawed and centrifuged at 13.2K rpm for 10 minutes. Protein (in water) was added in equal volumes to the buffer and titrant solutions with Hamilton syringes for a total concentration of 0.05 mg/mL. The buffer and titrant solutions, each with protein, were dispensed with a Microlab 600 dual syringe diluter pump (Hamilton Company) that was programmed to make a series of 16-20 samples at a total volume of 2 mL each. The set of samples contained different ratios of buffer:titrant solutions to achieve the final concentrations of denaturant of 0.2-4 M GdmCl, made at 0.2 M steps. Samples were incubated in a water bath at 25°C for one hour to reach equilibrium prior to taking fluorescence measurements. This is sufficient time for SNase to reach equilibrium in solution. The maximum time it takes for a protein folding reaction to come to equilibrium was determined from mixing two solutions to a final denaturant concentration that is the concentration midpoint in the transition region of the unfolding curve, as it is the farthest point from equilibrium. Buffer and titrant were combined at volumes to achieve the midpoint denaturant concentration and kinetic fluorescence measurements were recorded instantly and continually until the fluorescent signal stabilized. The time it took to reach equilibrium was consistently under 40 minutes.

Kinetic fluorescence measurements were collected for each sample with ATF-107 automated fluorometer (Aviv Inc.) in a quartz cuvette. The same excitation and emission parameters were used as in automated titration experiments. The voltage of the photon multiplier tube was fixed at the start of the experiment for the sample of protein in buffer and 0 M denaturant. The recorded fluorescence measurements were averaged for 10 seconds. Data were normalized and fit for the unfolding free energy and m-value in water per the same procedure as described for automated titration experiments in the main text.
